# Ectopic overexpression of abiotic stress-induced rice annexin, *OsAnn5* potentiates tolerance to abiotic stresses

**DOI:** 10.1101/2019.12.16.869131

**Authors:** Prasanna Boyidi, Trishla Vikas Shalibhadra, Halidev Krishna Botta, Deepanker Yadav, Pulugurtha Bharadwaja Kirti

**Author notes:** Corresponding authors at Department of Plant Sciences, School of Life Sciences, University of Hyderabad, Prof. C. R. Rao Road, Gachibowli, Hyderabad, 500046, India. E-mail: Prasanna Boyidi, P. B. Kirti.

## Abstract

The current study on putative rice annexin *OsAnn5* was tried to know its functional role in the abiotic stress tolerance. For this an *in silico* analysis of its protein sequence and upstream region was carried out. This results in identification of several probable potential sites for post-translational modifications and cis-elements respectively. We have studied the effect of *OsAnn5* in the amelioration of abiotic stress tolerance through heterologous expression in transgenic tobacco and *E*.*coli*. It is observed that *OsAnn5* over expression leads to enhanced tolerance to abiotic stress through efficient scavenging of the ROS and balanced expression of SOD and CAT antioxidant enzymes in both the systems, under stress treatments. Fluorescent signal for transiently expressed EGFP:OsANN5 fusion protein was localized in the peripheral region of the onion epidermal cells under salt stress treatment. Expression analysis of *OsAnn5* under ABA synthesis inhibitor, fluridone and salinity stress revealed that OsAnn5 appears to act through an ABA-independent pathway under salt stress and in support to this 35S:*OsAnn5* transgenics seedlings exhibited less sensitivity to externally applied ABA.

## Introduction

Annexins belong to a diverse multi-gene, multi-functional family of proteins (Moss et al., 2004; Clark et al., 2012, Yadav et al., 2018). They have been identified in fungi (Grenville-Briggs et al., 2010; Khalaj et al., 2015) plants, animals, aarchaea and also in prokaryotes (Kodavali et al., 2014). Annexin superfamily has been grouped into seven subfamilies, which are designated as ANNXA to ANNXG. Among them, plant annexins were grouped under the ANNXD family; however, all the annexins exhibit the conserved core domains in their protein structure (Moss & Morgan, 2004). Despite having highly conserved domains in their sequence, each annexin in a subfamily has been evolved for performing certain functions (Clark et al., 2012). They can bind the membrane in a Ca^+2^ dependent or independent manner (Dabitz et al., 2005). Variation in the expression level of annexins led to negative or positive effects on plant growth (Konopka-Postupolska et al., 2009; Zhu et al., 2014). Research on plant annexins revealed their involvement in growth and development (Chen et al., 2016; Huang et al., 2013; Proust et al., 1996; Zhu et al., 2014), amelioration of abiotic stress (Yadav et al., 2018) and biotic stress (Jami et al., 2008; Vandeputte et al., 2007). They are not only diversified in their tissue and sub-cellular localization (Baucher et al., 2012; Clark et al., 2000; Hoshino et al., 2004), but also in various functions (Buitink et al., 2006; Clark et al., 2001; Dai et al., 2006; Gallardo et al., 2003; Proust et al., 2005; Yang et al., 2007; Bassani et al., 2004; Clark et al., 2005; Sheffield et al., 2006; Yang et al., 2008; Bai et al., 2010). They are also known for their involvement in membrane binding, exocytosis (Battey et al., 1993), callose synthase activity (Andrawis et al., 1993), peroxidase activity (Gorecka et al., 2005, Jami et al. 2008), ion conductance (Laohavisit et al., 2009; Hofmann et al., 2000a), actin binding (Calvert et al., 1996; Huang et al., 2013; Hoshino et al., 2004a), ATPase activity (Mcclung et al., 1994) and GTPase activity (Shin et al., 1999). But, the exact signaling pathway they involved in and their mechanism of action are still largely unknown (Yadav et al. 2018).

Abscisic acid (ABA) is a key signaling hormone, which has been known to significantly affect plant metabolic pathways under abiotic stress conditions. Drought and salt stress signals lead to the accumulation of ABA which, in turn, regulates the expression of various stress related genes (Luchi et al., 2001; Shinozaki et al., 2000; Tuteja et al., 2007). Reports suggest that ABA can be a general regulator of annexins expression (Lee et al., 2004; Hoshino et al., 2004). Fluridone, an inhibitor of the carotenoid biosynthesis is known to inhibit ABA synthesis, which has been used earlier to decipher the regulation of genes under the control of ABA (Moreno et al., 2001; Toyofuku et al., 2000; Renouard et al., 2012; Barrero et al., 2008). In the present study, fluridone was used to identify whether the *OsAnn5* expression is ABA dependent or not during salt stress in its native system, rice.

Out of the ten rice annexins identified in the rice genome (Jami et al., 2012), OsANN1 has been characterized recently for its functional role in calcium-mediated heat stress tolerance in rice (Qiao et al., 2015). Knock out of another annexin, *OsAnn3* in rice plants lead to susceptibility under cold stress (Shen et al., 2017). However, the function of other annexins still needs to be ascertained. An earlier investigation on annexin gene expression has identified a rice annexin Os08g32970 (*OsAnn5*) to be highly expressive in several abiotic stress treatments (Jami et al. 2012). In this study, an *in silico* analysis was performed to understand the probable functional post-translational sites available in deduced protein sequence and potential *cis*-elements present in the putative promoter region of *OsAnn5*. Ectopic expression overexpression *OsANN5* in tobacco and in *E*.*coli* cells were performed to assess its involvement in abiotic stress, and the purified protein was assessed for its calcium binding activity. In the present investigation, we made an attempt towards the functional characterization of *OsAnn5* by over-expression in tobacco and tried to decipher its role in ABA signaling pathway in rice.

## Materials and methods

### In *silico* analysis of *OsAnn5*

Ten annexin CDS sequences were obtained from Rice Genome Annotation Project database (RGAP) (http://rice.plantbiology.msu.edu/) using the functional term search ‘annexin’. Out of these, Os08g32970 was selected and when this sequence was subjected to BLAST analysis, it showed high similarity to the *Ann5* of other plants species. Hence, it was named as *OsAnn5*. A 1.5 Kb upstream region of the Os08g32970 sequence was retrieved from RGAP using OsANN5 as a search query. The promoter sequence was analyzed through the online promoter analysis tool PLACE (http://www.dna.affrc.go.jp/PLACE/). The protein sequence was analyzed for post-translational modifications using various bio-informatics tools (Supplementary Table 2).

### Isolation and amplification of *OsAnn5*

Two weeks old *Oryza sativa* var *japonica* cv. Nipponbare (kindly provided by Dr. M. Seshu Madhav, Indian Rice Research Institute, Hyderabad) seedlings were subjected to 200 mM salt stress for 24 h and total RNA was isolated by RNAiso plus (*Takara*, China). Subsequently, cDNA was synthesized using M-MLV

Reverse Transcriptase (Sigma, USA). Full-length *OsAnn5* CDS was amplified by PCR using primers *OsAnn5* FP, *OsAnn5* RP (Supplementary Table 1) with the cDNA as the template. The amplicon was cloned in pTZ57R/T (Thermo Fischer Scientific Inc., USA) and the clone was confirmed by sequencing.

### Cloning and overexpression of *OsAnn5* in *E.coli* cells

The sequence confirmed *OsAnn5* was cloned into the pET32a+ bacterial expression vector (Novagen, USA) at the *Kpn*I and *Bam*HI sites so as to translationally fuse the C-terminal region of OsANN5 protein with 6x Histidine tag. The confirmed recombinant pET32a+:*OsAnn5* plasmid was transformed into *E*. *coli* BL21 (DE3) pLysS cells (Takara, China). The transformed BL21 cells were cultured in Luria broth at 37°C till 0.5 O.D growth and the cells were then induced with 0.1 mM IPTG (Sigma, USA) and grown further at 28°C for 3 h. The culture was then centrifuged at 5000 rpm for 10 min at 4^0^C. The pellet was resuspended in cold lysis buffer (50 mM ris–HCl, 300 mM NaCl, 10 mM imidazole and 0.1% N-lauroyl sarcosine, pH 8) an6d the cells were subjected to 20 s pulse of sonication and 20 s of pause for 25 cycles on ice. The lysed culture was centrifuged at 12,000 rpm for 40 min at 4^0^C. The clear supernatant was then gently mixed with 1 ml of Ni^+2^-NTA agarose beads (Qiagen, Germany) for 30 min and then added to the column with a flow rate of 0.5 ml/min. Later, the column was washed twice with the cold washing buffer (50 mM Tris–HCl, 300 mM NaCl and 20/30 mM imidazole, pH 8.0) to remove the impurities and the pure his tagged protein was eluted with the cold elution buffer (50 mM Tris–HCl, 300 mM NaCl and 200 mM imidazole pH 8.0) was confirmed as OsANN5 by SDS-PAGE (12%) and western blot using anti-His primary antibodies (Supplementary Fig. 1)

Along with the 6x Histidine tag, the thioredoxin (Trx) tag was also fused to OsANN5. Hence to cleave the Trx tag, thrombin enzyme was used. The cleavage reaction consisted of 0.01 units of thrombin enzyme (Novagen, USA), 1 X cleavage buffer and 100 µg of Trx:OsANN5 fusion protein in a final volume of 200 µl and was incubated at 16°C for 12 h. The tag cleaved OsANN5 was purified and dialyzed against 20 mM Tris-HCl. The calcium binding assay consisted of 200 µg OsANN5, 20 mM Tris, 50 mM NaCl and 2 mM CaCl_2_ at pH 6.0 in a final volume of 1 ml. Horiba Jobin Yvon florimax-3 fluorescence spectrophotometer was used for recording the fluorescence spectra in the near UV range of 200 nm to 400 nm.

### Ca^+2^ binding activity assay

The *OsAnn5* calcium binding activity assay performed in two methods. In the first method, three different concentrations of the CaCl_2_ i.e 10 mM, 20 mM, 30 mM CaCl_2_ was added to the OsANN5 overexpressed BL21 cell lysate samples to precipitate the calcium bound proteins (Qiao et al., 2015). Then each sample was subjected to centrifugation (12,000 rpm for 15 min) and the supernatant, precipitate were separated. Later, the respective precipitates were resuspended in the lysis buffer which has Na2EDTA (equal concentration to the added CaCl_2_) to chelate the Ca^2+^ ions. This resulted in the clearing of the precipitate. Subsequently, all the samples were analyzed through SDS-PAGE.

In the second method, fluorescence spectrophotometer (Horiba Jobin Yvon florimax-3) was used. The assay reaction consisted of 200 µg OsANN5, 20 mM Tris, pH 8.0 with or without 2 mM CaCl_2_ in a final volume of 1 ml. The fluorescence spectra were taken in the near UV range of 200 nm to 400 nm.

### Stress assays on *E.coli* cells

Spot assays were carried out to check the stress tolerance induced by OsANN5 in the bacterial system *E. coli*. For this, we followed the protocol used by Zhou et al., 2014. In brief pET32a::*OsAnn5* and pET32a+ transformed BL21 PLyS cell cultures were grown to appropriate cell density and induced as described above. Then different dilutions (10^-0^, 10^-1^, 10^-2^, 10^-3^, and 10^-4^) of both cultures were spotted and air dried on LA media culture plates (100 mg/L ampicillin and 0.1 mM IPTG) containing various concentrations of salt, 0.4 M, 0.5 M, 0.6 M NaCl (high salinity stress), 0.8 M sorbitol (osmotic stress) and 8% and 10% PEG (drought stress). All the plates were containing ampicillin as selection agent. For the high temperature stress treatment after 3 h of IPTG induction, the culture was grown at 50 ^0^C for 30 min, and 1 h. The respective dilutions were spotted on Luria Agar selection plates. The plates were incubated at 37°C overnight and checked for their growth performance.

### Generation of *OsAnn5* tobacco transgenic plants

The *OsAnn5* was cloned into pRT100 vector (Töpfer et al., 1987) using *KpnI* and *BamHI* restriction enzyme sites for a transcriptional fusion with the CaMV 35S promoter and polyadenylation signal. This expression cassette CaMV35S:OsAnn5:polyA was cloned into pCAMBIA2300 binary vector (CAMBIA, Australia) at the *Hin*dIII restriction enzyme site and the recombinant vector was confirmed by digestion and PCR (Supplementary Fig. 2). This recombinant pCAMBIA2300 binary vector carries *nptII* as a plant selection marker and was mobilized into Agrobacterium strain LBA4404 for developing tobacco transgenics.

Ectopically expressing *OsAnn5* tobacco transgenic plants were developed using the standard leaf disc transformation protocol developed by Horsch et al., (1985). In brief, leaves from *in vitro* grown two weeks old *Nicotiana tabacum* cv Samsun plants were used as explants. They were cut into small pieces and co-cultivated with *Agrobacterium tumefaciens* strain LB4404 carrying the recombinant binary vector on MS medium having 2.0 mg/L BAP and 0.1 NAA mg/L in dark for 48h. The explants were then subjected to shoot regeneration on MS medium supplemented with 2.0 mg/L BAP, 0.1 NAA mg/L, 125 mg/L kanamycin and 250 mg/L Cefotaxime under 16/8 h light-dark photoperiod. Regenerated shoots were transferred to the rooting medium (0.5x MS salts, 0.1 mg/L NAA and 125 mg/L kanamycin). Rooted T0 plants were transferred to sterile soil in small cups for acclimatization and well established plants were shifted to pots and allowed to set seeds by self-pollination in the green house.

### Confirmation of 35S:*OsAnn5* tobacco transgenic plants

The putative primary transformants were confirmed by PCR for the presence of *OsAnn5* using specific primers for *OsAnn5* and *nptII* genes. A total of 15 independent 35S:*OsAnn5* transgenics plants were developed. The PCR positive primary transgenic plants were analyzed for low and high expression using a semi-quantitative PCR. The seeds from putative primary transgenic plants were germinated on a half strength MS medium supplemented with 135 mg/l kanamycin to raise the T_1_ progeny and to identify the null segregants. The null segregants (NS) were later rescued on MS medium without the selection agent and grown to maturity for use as controls in the stress experiments. The T_2_ generation homozygous lines were obtained by germinating the self-pollinated seed of T_1_ plants on antibiotic selection and identification of 100% germination of the progeny seedlings on the kanamycin selection medium. All the stress assays were conducted with either T_2_ seeds or plants.

### Seed germination and seedling assay

For seed germination assay, approximately 100 seeds of each transgenic line and the Null Segregant control (NS) were surface sterilized with 2% sodium hypochlorite for 15 min and then washed thrice with autoclaved distilled water, placed on half strength MS media as a control and half strength media containing 200 mM NaCl. The seed germination was monitored every day and observations were recorded.

For the seedling assay, ten days old NS and transgenic seedlings were placed on the different 0.5X MS media containing 200 mM NaCl, 0.8 M sorbitol, and 10 % PEG. After 20 days of treatment, the phenotype was observed and their root length was calculated for comparison studies.

### Root length assay

10 days old transgenic and NS seedlings were kept on stress media and grown for 2 weeks. After the treatment the seedlings were arranged in equally heights and their root length was compared, and measured. The root length was calculated and expressed in fold difference.

### Lea disc assay

Six weeks old NS and transgenic plants grown in the green house were used for leaf disc assay. Fully grown mature leaf from the plant was taken and one cm diameter leaf discs were cut in water and subjected to different abiotic stresses. After one week of incubation, their phenotype was observed and the samples were analyzed for total chlorophyll content, proline and MDA levels.

### Chlorophyll estimation

The total chlorophyll content was estimated using the protocol detailed out by Hiscox et al. (1979) with minor modifications. All the stress treated and control samples (50 mg) were extracted in 80% acetone on ice and the extract was centrifuged at 10,000 rpm for 10 min at 4 ^0^C. The absorbance was recorded for the supernatant at 645 and 663 nm in a spectrophotometer (SHIMADZU, spectrophotometer, UV-1800). Total chlorophyll content was calculated using Arnon’s equation and the values were expressed in μg/g FW.

Total chlorophyll (mg/gFW) = 20.2 (A645) + 8.02 (A663)]*V/1000*W

### Proline estimation

Proline content was estimated through a standard protocol described by Bates et al. (1973) with minor modifications. The leaf samples (50 mg) of control and stress treated NS and transgenic plants were homogenized in 1 ml of 3% of sulfosalicylic acid. The homogenate was centrifuged at 10,000 rpm at 4°C. One hundred μl of supernatant was added to 200 μl glacial acetic acid and 200 μl acid ninhydrin mixture. The total reaction mixture was boiled at 100°C for 1 h. After incubation, the reaction was stopped by placing the samples on ice and equal volumes of toluene were added to each sample until proline was extracted into the inorganic phase from the aqueous phase. The absorbance of the extracted samples was measured at 520 nm using toluene as a blank. Based on the standard curve, proline concentration was determined and expressed as μg g^-1^ FW.

### Lipid peroxidation assay

Lipid peroxidation levels were estimated through the melondialdehyde (MDA) quantification (Heath et al., 1968). Each sample (50 mg) was homogenized in 1 ml of 0.1% TCA and the homogenate was centrifuged at 12,000 rpm at 4°C for 10 min. Subsequently, 500 µl of the supernatant was mixed with 1.5 ml of 0.5% (w/v) TBA, which was prepared in 20% TCA (w/v) and incubated at 95°C for 30 min. The reaction was stopped by placing the tubes on ice followed by centrifugation for 5 min at 12,000 rpm (4°C). The absorbance values for the supernatant were taken at 532 and 600 nm. MDA concentration was calculated using its molar extinction coefficient (155 mM^-1^cm^-1^). Results were represented as µmol/gFW. The formula used was: MDA (mM) = (A532 - A600)/155

### Hydrogen peroxide detection by DAB (3, 3′-Diaminobenzidine) staining and quantification

Diaminobenzidibe (DAB) staining was performed to detect the formation of hydrogen peroxide (H_2_O_2_) in the stress treated samples according to the protocol developed by Daudi et al. (2012). In brief, the fourth leaf from one month old transgenic and NS plants was taken for the analysis. Leaf discs were treated with 250 mM NaCl for 48 h. After the treatment, they were incubated in DAB staining solution (1 mg/ ml DAB, pH 7.4) overnight. Later, the chlorophyll was removed from the samples by boiling in bleaching solution (ethanol: acetic acid: glycerol = 3:1:1) for 20 min. Subsequently, they were stored in 10% glycerol until images were captured.

The DAB stained leaf discs were homogenized in liquid Nitrogen and extracted in 1 ml of perchloric acid and centrifuged at 10,000 rpm for 10 min. The absorbance for the supernatant was taken at 450 nm. The H_2_O_2_ levels were represented in the graph as µmole/gFW.

### Antioxidant enzymatic assays in *E. coli* and tobacco

The pET32a (V) and pET32a:*OsAnn5* (G) transformed bacterial cells were cultured in Luria Broth containing 0.6 M NaCl as salinity stress with or without induction. For the heat stress, the induced and un-induced cultures of V and G were subjected to 50 °C for 30 min. All the samples (vector (V) un-induced or induced (VI), gene carrying vector (G) un-induced or induced (GI), vector salt treated un-induced (VS) or induced (VSI), gene carrying vector salt treated un-induced (GS) or induced (GSI), vector heat treated un-induced (VH) or induced cultures) were centrifuged at 5,000 rpm for 10 min and frozen in liquid nitrogen (N2). Later, they were washed with PBS, resuspended in the 50 mM PBS (pH 7.8) and subjected to sonication. After sonication, the crude lysate was centrifuged at 12,000 rpm for 30 min and the clear supernatant was used for enzymatic assays. Un-induced cultures for control, high salinity and heat stress were taken as controls.

For tobacco, the control and 200 mM NaCl treated leaf samples of NS and 35:OsAnn5 transgenic plants were frozen in liquid N_2_ and homogenized in 50 mM phosphate buffer (pH7.8). Further, the same samples were centrifuged at 12,000 rpm for 30 min and the supernatant was used for superoxide dismutase (SOD) and catalse (CAT) enzyme assays.

The protein concentration was estimated through Bradford reagent using BSA standard curve. SOD activity assay was performed according to the protocol developed by Beyer and Fridovich, (1987) with minor modifications. The reaction mixture for SOD activity contained 0.66 mM Na2EDTA, 10 mM L-methionine, 33 µM NBT and 0.0033 mM riboflavin and 100 µg of protein in 50 mM sodium phosphate buffer (pH7.8). The final reaction mixture was kept in light for 15 min. Later the blue color intensity was measured at 560 nm. One unit of SOD activity is defined as the amount of protein required to inhibit the photo-reduction of NBT by 50 %. The final activity of all samples was represented in units/mg/ gFW.

CAT activity was measured using the protocol developed by Çelik & Atak, (2012) with little modifications. The CAT enzymatic reaction mixture includes 25 µg of protein, 50 mM phosphate buffer (pH7.0) and 19.8 mM H_2_O_2_. The decreased absorbance for H_2_O_2_ was taken at 240 nm. Units of activity were calculated using the H_2_O_2_ *molar extinction coefficient* (43.6 M^-1^ c.m^-1^) and the final activities were expressed as units/mg/min.

### Transient expression of EGFP:*OsAnn5* in onion epidermal peels

*The OsAnn5* sequence was translationally fused to the N-terminal of EGFP in the pEGAD vector (Cutler et al., 2000) at the *Eco*RI and *Bam*HI restriction sites. The confirmed EGFP:*OsAnn5* construct was used for transient expression in the onion epidermal cells through *A. tumefaciens* mediated transformation protocol as described by Sun et al. (2007). In brief pEGAD:EGFP:*OsAnn5* transformed agrobacterium culture was grown at 28°C for overnight under the rifampicin (50mg/L) and kanamycin (100 mg/L) selection. Later the culture was pelleted down, resuspended in the MS medium (100 µM aceto syringin, 5 % sucrose, 0.02 % silwet-77) and incubated for 6 h. Then fresh onion scales (1 cm x 1 cm) were placed in the agrobacterium resuspension solution for 6 h as their inner surface immersed in it for infection. Later the infected onion scales were transferred to co-cultivation media (½ MS) for 2 days. After co-cultivation onion scales were rinsed with 0.1 M phosphate buffer and their epidermal peels were peeled off and kept on the glass slide to check under the confocal microscope. Fluorescence images were taken under 20X lens.

### Quantitative Real-time PCR

To check the role of *OsAnn5* in ABA signaling pathway, 15 d old *Oryza sativa* var *japonica* cv. Nipponbare seedlings were subjected to different treatments viz, 100 µM ABA, 200 mM NaCl, 100 µM fluridone (ABA synthesis inhibitor) and 200 mM NaCl+100 µM fluridone. The samples were collected at different time points i.e. 6, 12, 24 h. Before treating the rice seedlings with 200 mM salt + fluridone, they were pre-treated with 100 µM fluridone for 12 h. After the treatment, all the samples were subjected to RNA isolation followed by cDNA synthesis. Subsequently, real-time PCR was performed to analyze the gene expression levels.

Quantitative real-time PCR was performed with the Bio SYBR Premix Ex Taq (Takara, China) according to the manufacturer’s protocol. 100 ng of cDNA was used as a template for the reaction. The reaction was set up in Axygen 96-well PCR microplates with the following reaction conditions: initial denaturation at 95°C for 30 sec, 40 cycles of 95°C for 3 sec, 58°C for 30 sec. Raw data was analyzed through the ΔΔC_T_ method (Livak et al., 2001). Sigma Plot 11 was used for the representation of the data.

### Statistical analysis

All the experiments conducted in this study were repeated thrice and the values plotted are the average of three independent experiments. The ssignificance values were calculated using one way ANOVA in the SIGMA PLOT 11. * *P<* 0.05, * * *P<* 0.01, * * * *P*< 0.001.

## Results

### *In silico* analysis of promoter and protein sequences of *OsAnn5* reveals the presence of potential *cis*-elements and functional motifs

*In Silico* analysis of the promoter sequence showed the presence of various putative *cis*-acting elements known to be involved in abiotic stress responses like dehydration, low temperature and salt stresses were observed in the putative promoter region. Important motifs for hormonal response for auxin, gibberellic acid, ABA, ethylene and cytokinin, and light response were observed in the amino acid sequences. Tissue-specific elements for mesophyll, pollen, root hair-specific expression were also observed in the putative promoter region. The binding sites for various transcription factors such as MYB, MYC, WRKY and DOF were recorded in the promoter region. Several light responsive elements were also found in it (supplementary table 3). This data suggests that OsANN5 is highly regulated during growth and plays a major role during stress responses. This also indicates that *OsAnn5* expression is regulated during different environmental and abiotic stress conditions and is consistent with the *cis*-acting elements predicted in its putative promoter sequence

Post translational modification (PTM) is another kind of regulatory mechanism, which expands the functional range of the proteins. Various bioinformatics tools were used to identify the sites for possible PTMs in OsANN5 protein. This analysis resulted in the identification of some potential phosphorylation sites and disordered regions in the protein. Three disordered regions were identified in OsANN5, which spanned the amino acids at the positions, 2 – 15, 128 – 140, 314 – 319. Probable phosphorylated amino acids and the related kinases were identified as 44 S- Unspecified (Unsp) and PKC, 146 S- Unsp, PKA, 200 S- Unsp, 275 T- Unsp, PKC, 296 S- Unsp. Only those having prediction score of ≥ 0.9 were chosen for presentation. Several Lysine residues in the protein were possible residues that can be ubiquitinated and these amino acids were also predicted to have equal probability of being acetylated, viz., K 25, K 74, K 227, K 250, K 254, K 261, and K 294. We observed one potential site for N-Glycosylation with highest score at N 266. However, no sites for O-Glycosylation were observed in the amino acid sequence. Two consensus sequences for SUMOylation (K 227, K 286) and one SUMO interaction site (IRVVTT) were also identified. One calcium binding motif [XGT(38 residues)D/E] was identified in the protein. Interestingly, a stretch of six glycine residues (ARF**GGGGGGG**LEH) was observed in the protein sequence, which is not found in other rice annexins. As in other annexins, a conserved histidine residue,at the 40^th^ position and a motif for the salt bridge (Laohavisit et al., 2011), F-actin binding site have also been observed in the amino acid sequence (**Fig. 1**). Possible interacting partners for OsANN5 were identified as a hypothetical protein (LOC_Os06g34710.1) and a MYB family transcription factor (LOC_Os02g34630.1).

**Fig. 1.**
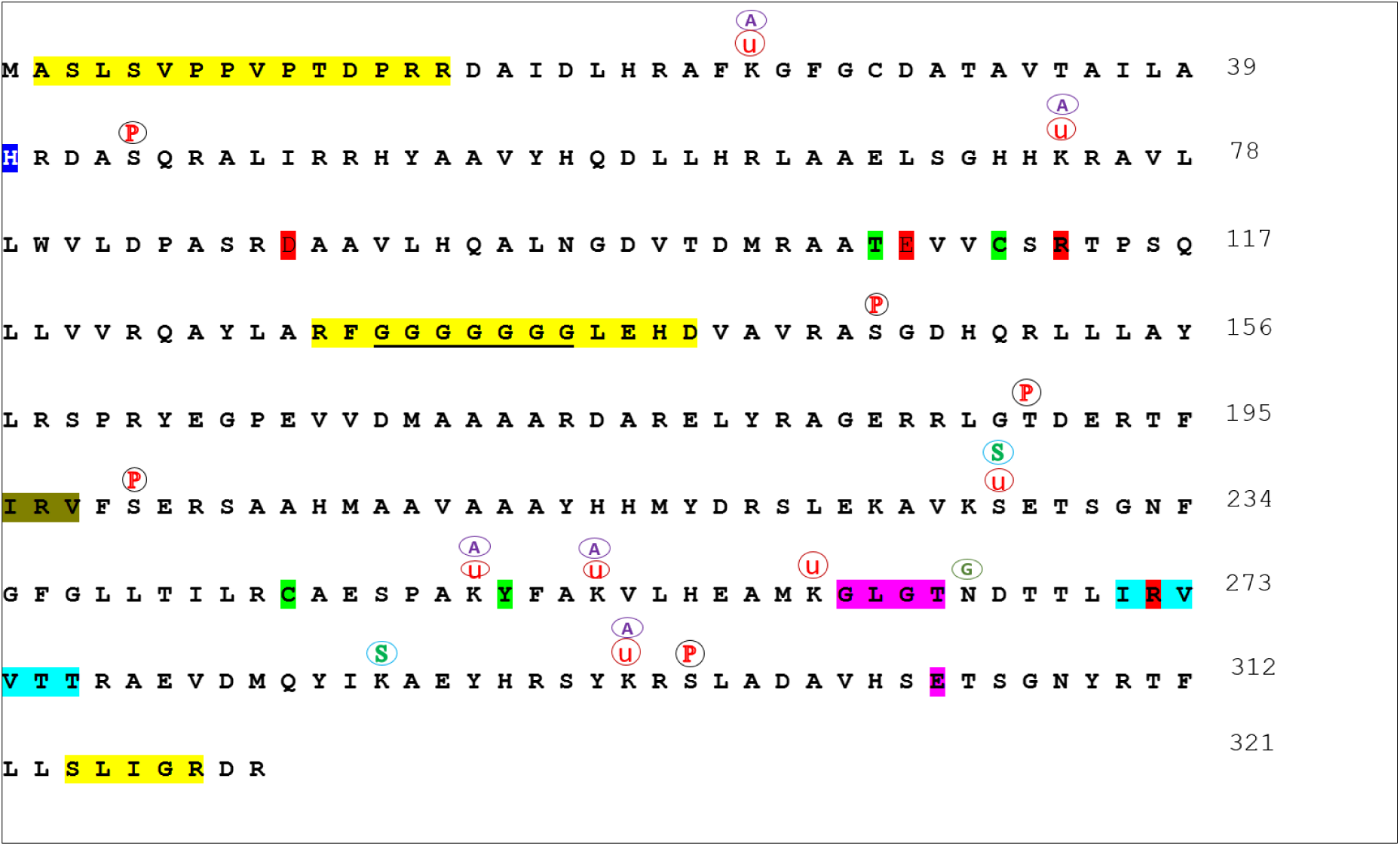
Mapping of putative Phosphorylation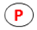, N-Glycosylation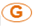, Sumoylation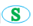, Ubiquitination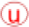 and acetylation sites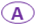 in OsANN5 sequence. Residues involved in salt bridges are in red, conserved histidine residue is highlighted in blue and disordered regions are highlighted in yellow, sumo interaction site is indicated in teal color. Underlined region is the glycine stretch, neon green highlighted amino acid residues are of S3 cluster, olive green highlighted region is the F-actin binding motif, pink highlighted region is the calcium binding motif (Endonexin fold)

### OsANN5 expression helps *E. coli* cells to combat abiotic stress conditions

To check whether the overexpression of OsANN5 improves abiotic stress tolerance in *E. coli,* 10µl of both pET32a transformed and pET32a:OsANN5 transformed BL21 pLysS cultures of different dilutions were spotted on Luria Agar media, which induce high salinity, osmotic, drought stresses along with the ampicillin selection. Their growth performance was observed after 16 h of incubation at 37 °C. Under all the abiotic stress conditions, *E. coli* cells overexpressing OsANN5 were grown well compared to the cells with vector transformation. High salt and heat stress shown major effect on the growth of the vector transformed cells. Under the salt stress the OsANN5 overexpressing *E*.*coli* cells took less time to adopt and grow. Similar tolerance was observed to the heat treatment (**Fig. 2**). After one hour heat stress (50 °C) treatment, vector cells did not recovered, but OsANN5 overexpressing cells were able to resume normal growth. This clearly suggests that the expression of OsANN5 helps the bacterial cells in combat the the abiotic stresses, especially to the salt and heat stresses.

**Fig. 2.**
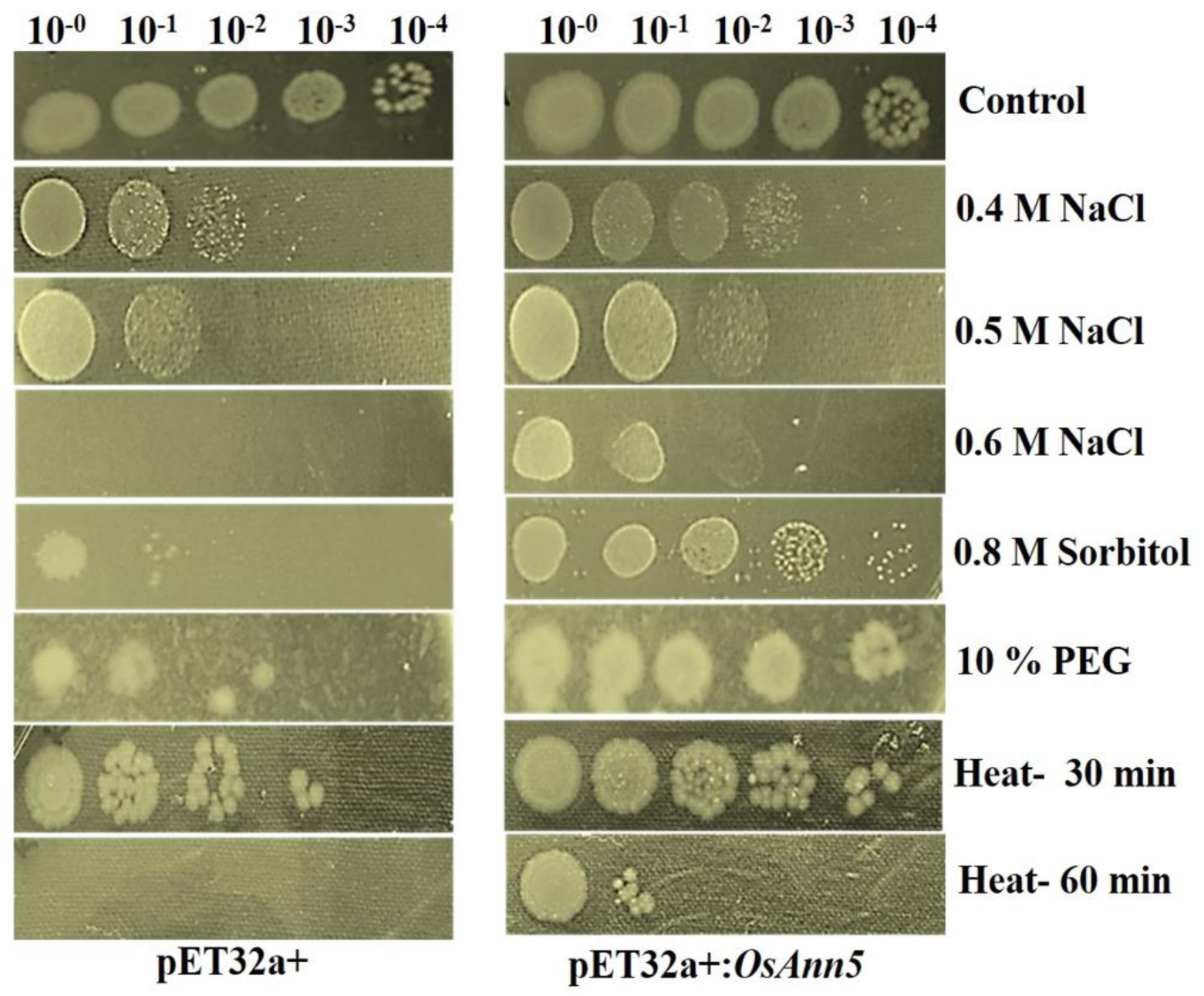
The pET32a+ and pET32a+:*OsAnn5* transformed culture were induced with 0.1 mM IPTG for 3 h at 28 °C. A 10 µl sample from different dilutions (10^-0^, 10^-1^, 10^-2^, 10^-3^, 10^-4^ ) of the both induced cultures were spotted on different LA plates (Ampicillin + IPTG) which contains 400 mM NaCl, 500 mM NaCl, 600 mM NaCl, 0.8 M sorbitol, 10 % PEG to induce the abiotic stress. The heat stress (50° C for 30 min and 1 h) given cultures were spotted on normal LA medium (100 mg/L Ampicillin + 0.1 mM IPTG). The spotted plates were incubated at 37 °C for minimum of 12-16 h and their growth was observed.

### Calcium binding activity of OsANN5

Most of the annexins are Ca^+2^ binding proteins but, not all of them (Dabitz et al., 2005) . To check the OsANN5 calcium binding ability there different concentration of CaCl_2_ i.e 10mM, 20 mM, 30 mM was added to the OsANN5 expressing *E.coli* cell lysate which led to the precipitation of the calcium binding proteins. SDS-PAGE was carried out to find out the proteins present in the precipitate and supernatant (**Fig. 3A**). Trx tag fused OsANN5 (50 KDa) was found in the precipitated proteins, not in the supernatant proteins. At 10 mM CaCl_2_ concentration less amount of OsAnn5 got precipitated. The 20 and 30 mM CaCl_2_ treatments leads to more OsAnn5 precipitation. This results imply that OsANN5 is a calcium binding protein.

**Fig. 3.**
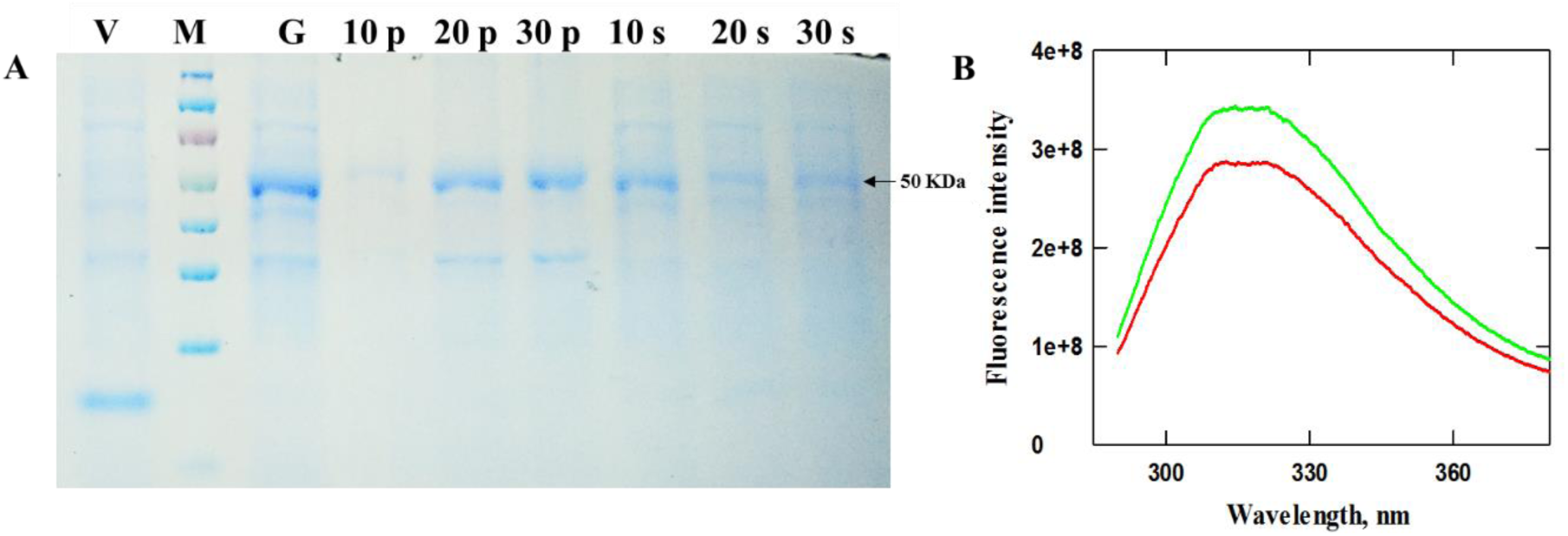
Ca^+2^ binding activity assay for OsANN5 through CaCl_2_ precipitation and fluorescence spectroscopy method. The IPTG induced pET32a+:*OsAnn5* transformed cell lysate was treated with different concentration of CaCl_2_ **i.e** 10 mM, 20 mM, 30 mM **(A).** This led to the precipitation of the calcium bound proteins (10p, 20p, 30p are precipitate proteins after 10mM, 20 mM, 30 mM CaCl_2_ treatment respectively) . After centrifugation the precipitated proteins were resuspended in the lysis buffer having the Na_2_EDTA and the supernatant having Ca^+2^ unbound proteins (10s, 20s, 30s are Ca^+2^ unbound proteins after 10mM, 20 mM, 30 mM CaCl_2_ treatment respectively) were loaded on the SDS-PAGE. V= pET32a+ transformed cell lysate, M=Marker, G= OsANN5 overexpressing cell lysate. Fluorescence emission spectra for OsANN5 with (Red) and without addition of 2 mM CaCl_2_ (green) **(B)**.

A Fluorescence Spectroscopy study was also conducted to assess the calcium binding activity of the OsANN5. Fluorescence emission for purified OsANN5 was detected at 320 nm wavelength. Upon addition of 2 mM CaCl_2_, a decrease in the fluorescence peak intensity was observed (**Fig. 3B**), suggesting that OsANN5 binds to Ca^+2^ ions, which led to fluorescence quenching and ultimately resulted in a shift in the fluorescence emission intensity. Insignificant or very less fluorescence emission was observed when an uncleaved OsANN5 (Trx:OsANN5) was used for the assay (data not shown).

### OsANN5 localizes in the peripheral under salt stress

Transient expression of EGFP fused OsANN5 was carried out in onion epidermal peels to detect its sub-cellular localization. We observed that the fluorescence signals were localized in the corners of the cells under normal condition. But, upon 200 mM NaCl treatment for 10 min, these signals were observed in the peripheral regions of the cells, but not in the cell wall (**Fig. 4**). In the vector transformed cells, the signal is diffused throughout the cell including nucleus.

**Fig. 4.**
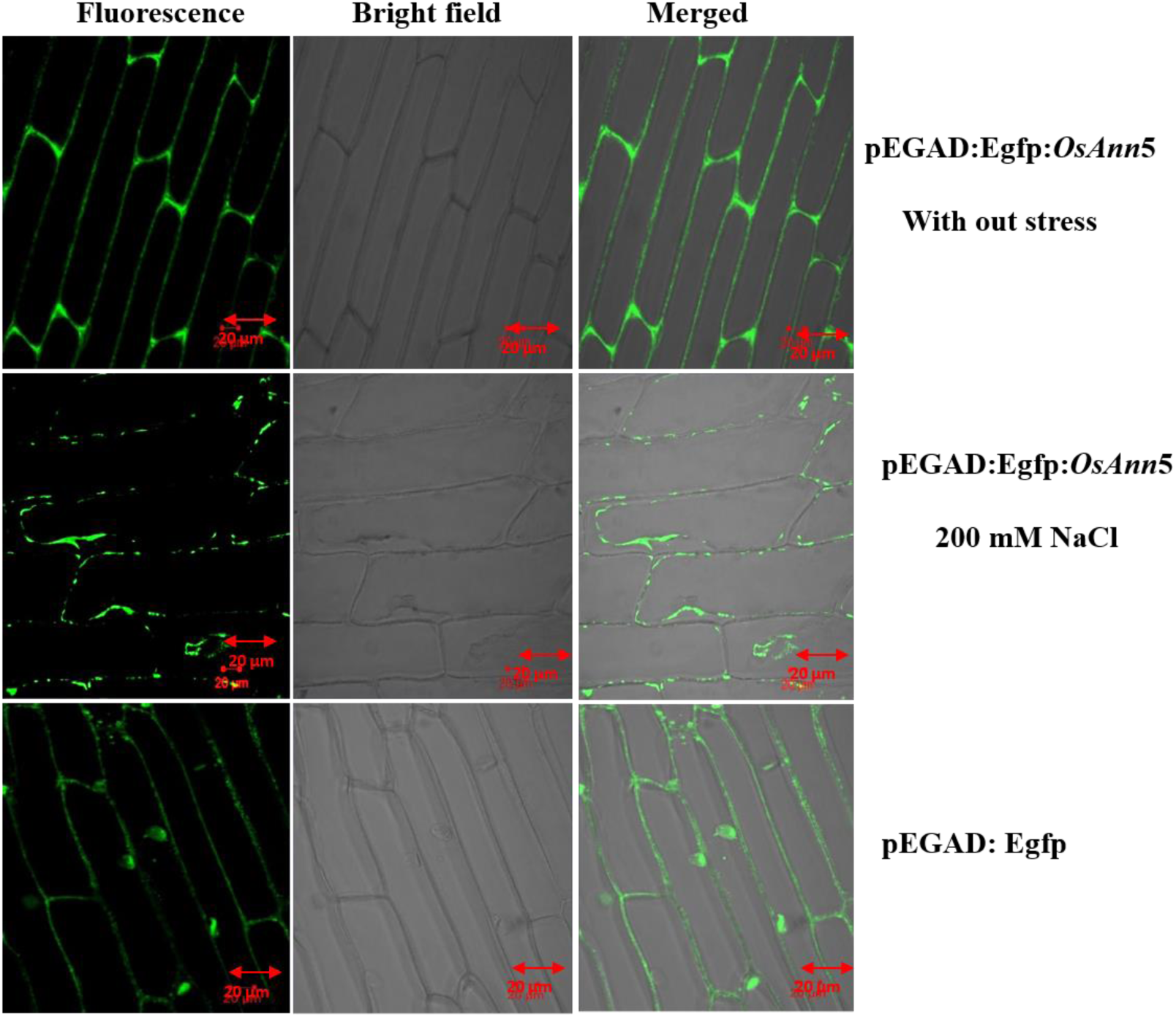
Subcellular localization studies on OsANN5 (**B**). Transient expression of EGFP fused *OsANN5* was done in onion epidermal peels. The fluorescence for Egfp fused *OsAnn5* transformed onion epidermal peels was observed in the corners of the cells. After NaCl treatment the fluorescence was observed in the peripheral region of the cell. In the vector (pEGAD::EGFP) transformed onion epidermal peels, the fluorescence was throughout the cell. Confocal images were taken with the 20X magnification with scale bar of 20 µm.

### Confirmation of 35S:OsAnn5 transgenics

The putative *35S*:*OsAnn5* tobacco transgenic plants were screened for the presence of *OsAnn5* and *nptII* in their genomic DNA (Supplementary Fig. 3). The confirmed transgenic plants were screened further for low and high expression of the target gene through Semi quantitative PCR analysis using ubiquitin as a reference gene (Supplementary Fig. 3). Of them 35S*:OsAnn5* line l-2 (A-2) was selected for low expression, while 35S:*OsAnn5* line- A4 (A-4) and 35S:*OsAnn5* line - A7 (A-7) were used to represent high expression for the stress assays. Various abiotic stress assays were carried out to assess the stress tolerance property of T3 35S:*OsAnn5* tobacco transgenics.

### Constitutively overexpressing *OsAnn5* tobacco transgenic shown tolerance to abiotic stress treatments

To assess the stress tolerance of 35S:*OsAnn5* transgenic plants, various stress assays like seed germination percentage, seedling assay and leaf disc assay were performed. Seed germination is an important factor affected by the abiotic stresses. This assay revealed that NS seeds took 10 d for 50 % of germination while the 35S:OsAnn5 transgenic plants showed 50% germination by the 8^th^ day (Fig. 5). Without any stress treatment, under control conditions all the seeds germinated by the 3^rd^ day.

**Fig. 5.**
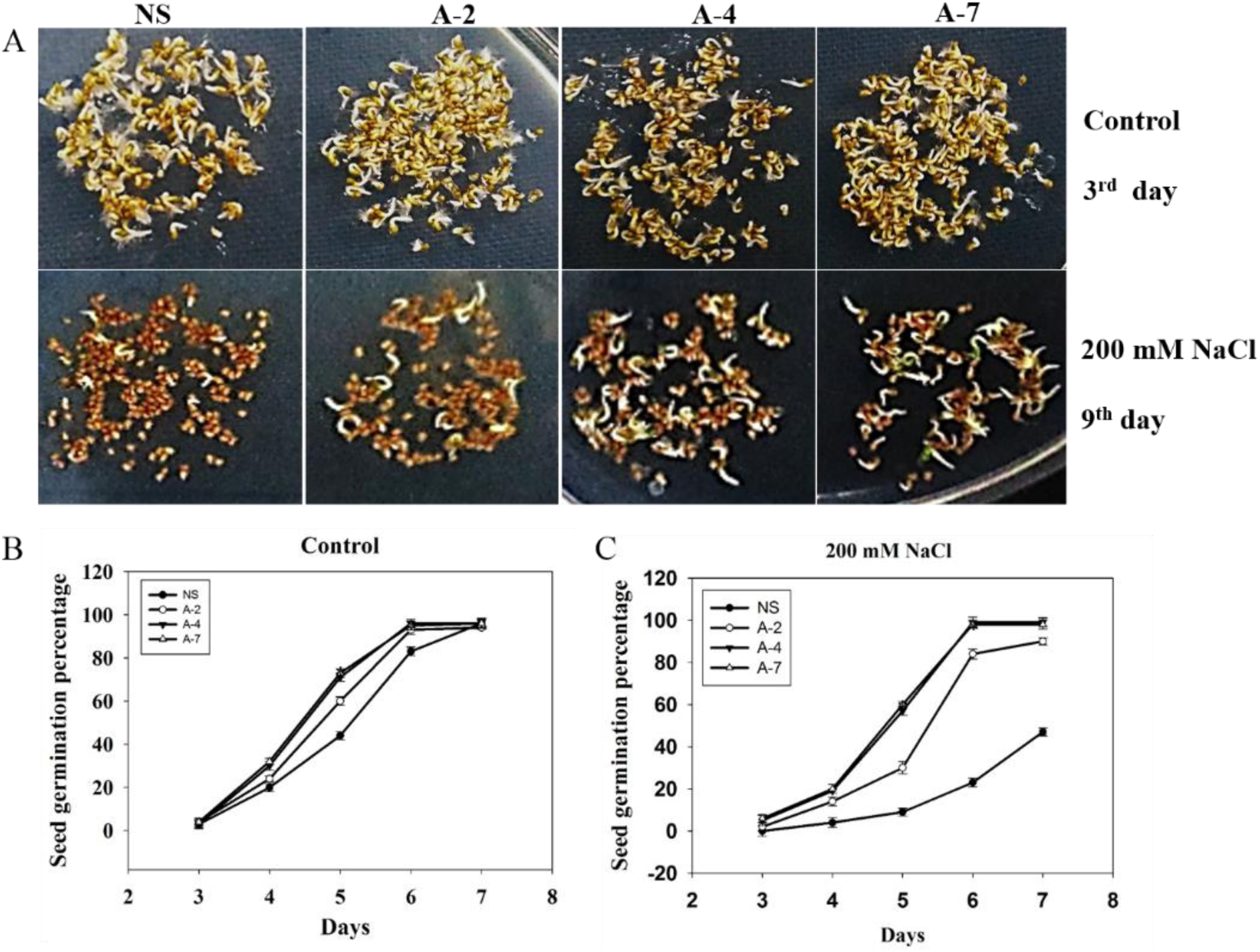
Seed germination assay on null segregants (NS) and 35S:*OsAnn5* transgenic plants. Approximately 100 seeds of NS and transgenics were placed on the ½ MS as a control and on ½ MS+200 mM NaCl media. The germinated seeds were counted on daily basis **(A).** Germination percentage under control **(B)** and NaCl **(C)** conditions were calculated and plotted in a graph. Asterisc represents the significant difference in the germination percentage in comparison with the NS when analyzed through ANOVA. * *P<* 0.05

For the seedling assay, NS and 35S:OsAnn5 T2 transgenic seeds were germinated on normal MS medium and kanamycin selection media respectively. After 10 d of germination and growth, the seedlings were subjected to abiotic stress treatments like high salinity (200 mM NaCl), drought (10% PEG) and osmotic stresses (300 mM Sorbitol) and the seedling responses were assessed to check their stress tolerance (**Supplementary Fig. 4**). Compared NS seedlings, the 35S:OsAnn5 transgenic seedlings were able to two fold reduction in the root length was observed for NS under all abiotic stresses while the 35S:OsAnn5 transgenic seedlings did not shown any significant reduction in their root length (Fig. 6).

**Fig. 6.**
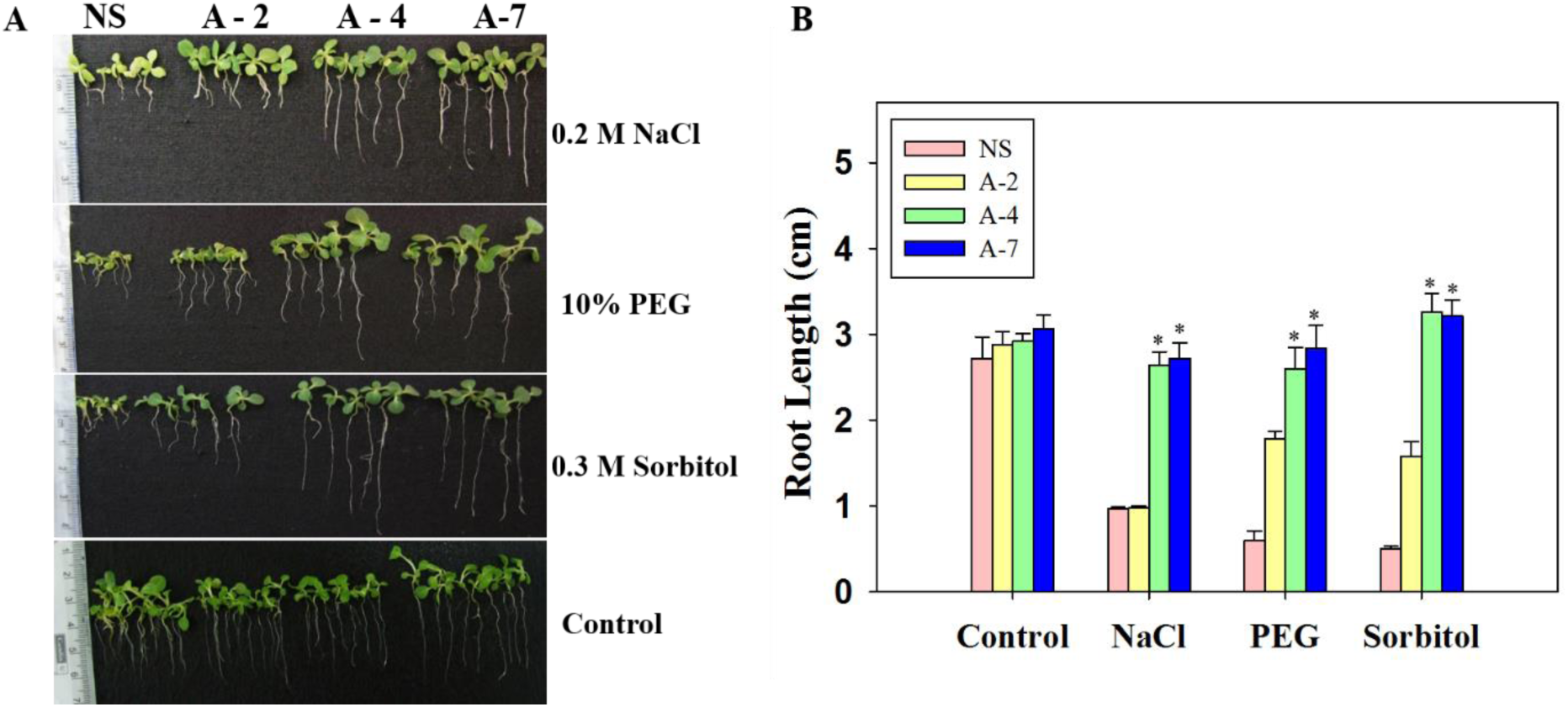
Abiotic stress assays on 10 days old NS and 35S:*OsAnn5* tobacco transgenics. The seedlings were grown on MS media supplemented with 200 mM NaCl, 10 % PEG, 0.8 M sorbitol and on MS media as a control. The difference in the root lengths after the treatment were compared and measured using the scale **(A)** and the values were represented in bar graph **(B)**. Error bars represent the ± SD. Asterisks represent the significant difference in the root length of null segregants and transgenics under stress conditions analyzed through ANOVA. \**P<* 0.05.

To check the extent of tolerance to the various abiotic stresses at mature plant stage, a leaf disc assay was performed for all transgenics and NS. The leaf discs were subjected to similar stress conditions and their phenotype was assessed (Fig. 7A). Subsequently, the samples were analyzed for their total chlorophyll content, proline content and the extent of lipid peroxidation under respective treatments.

**Fig. 7.**
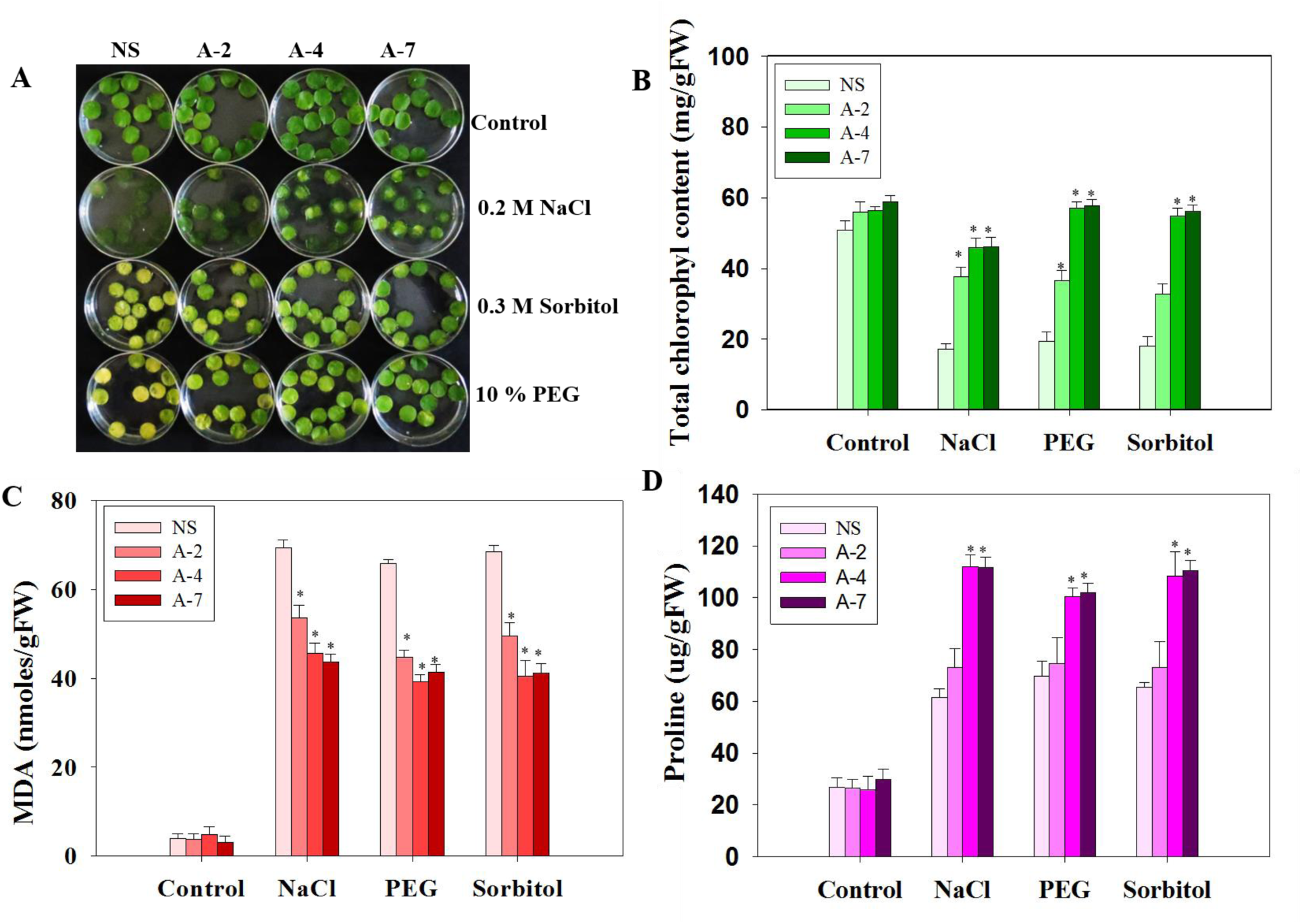
Leaf disc assay for null Segregant and 35S:*OsAnn5* transgenic lines under control and different abiotic stress (200 mM NaCl, 0.8 M Sorbitol, 10% PEG) conditions (A). After 7 d of the treatment the leaf disc samples were analyzed for total chlorophyll content (B), MDA levels (C), proline accumulation (D). Error bars represent the ± SD. Asterisks represent the significant difference in either fold level or percentage level in comparison with stress treated null segregants under respective stress conditions (NS) analyzed through ANOVA. * *P<* 0.05; ** *P<* 0.01.

The NS showed up to 60% reduction in the chlorophyll content under all the stress conditions whereas *35S*:OsAnn5 transgenic plants showed only 15 - 30% reduction (Fig. 7B). Proline accumulates in the plants during stress conditions and acts as a compatible solute, which in turns helps in protecting enzymes and proteins during stress (Szabados et al., 2010). The accumulation of free proline by the NS was one fold higher under stress as compared to the control condition. The transgenic tobacco plants accumulated significantly higher amount of free proline under the same stress conditions (Fig. 7D). The stress on the plant also intensifies lipid peroxidation, which affects the membrane integrity and hence, the viability of cell. MDA is the end product of lipid peroxidation process that represents the oxidation status of the cells during stress conditions. Significantly less MDA levels were present in the *OsAnn5* transgenic plants in comparison with their corresponding NS (Fig. 7C).

A DAB staining assay was performed to observe the extent of H_2_O_2_ accumulation under stress in the leaf discs (200 mM NaCl treatment). We observed that salt stress treatment lead to strong and intense staining of NS leaf discs when compared to the transgenic plants (Fig. 8B). DAB quantification results suggested that *OsAnn5* transgenics shown significantly less H_2_O_2_ accumulation under the salt stress compared to NS (Fig. 8B). These results suggest that NS were not able to scavenge the H_2_O_2_ as efficiently as the 35S:OsAnn5 transgenic plants.

**Fig. 8.**
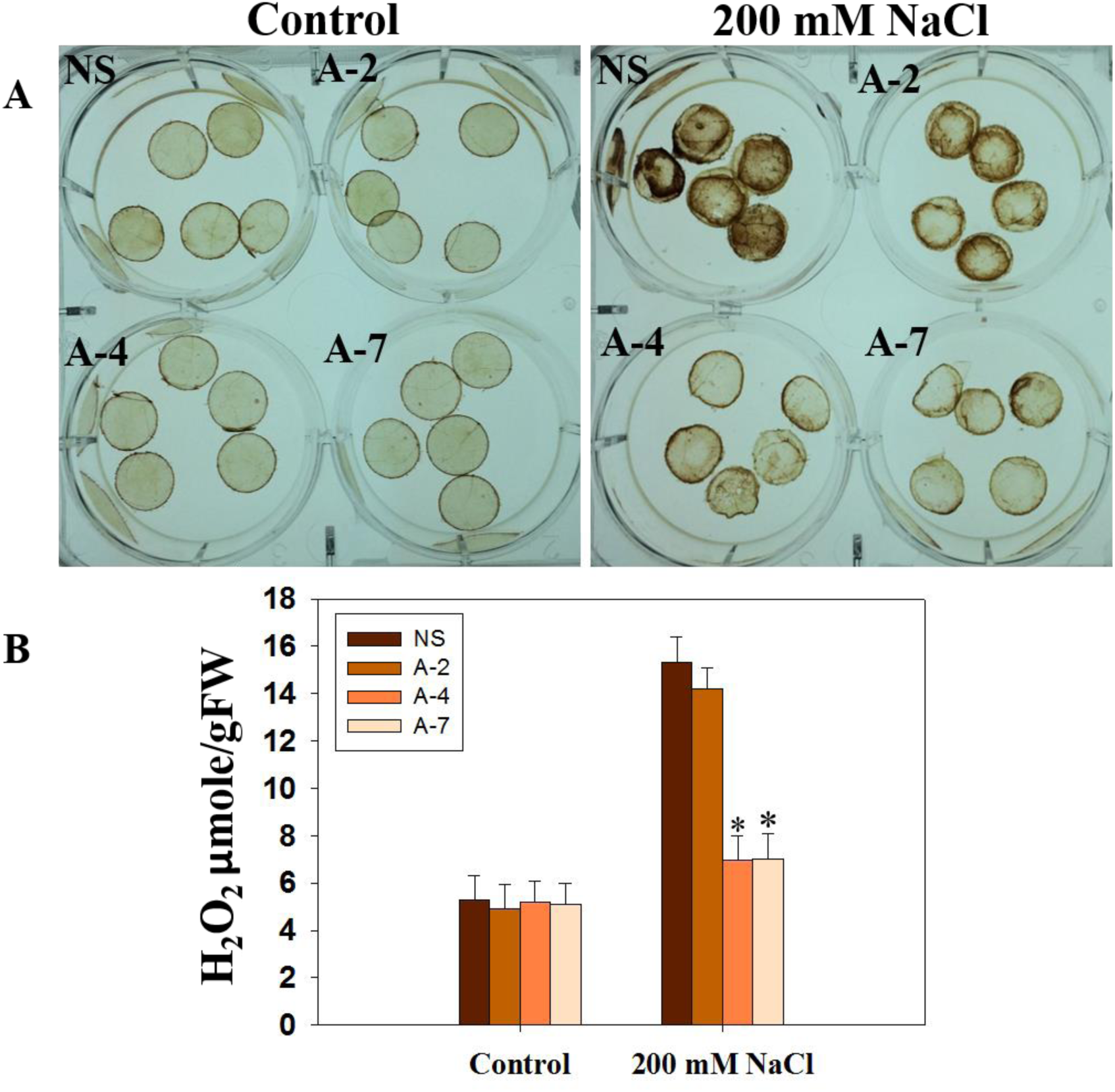
DAB staining assay was performed for the NS and 35S:*OsAnn5* transgenic lines. The brown coloration was directly proportional to the levels of H_2_O_2_ accumulation. The stress led to more intense staining of the NS leaf disc than transgenic lines (**A**). The DAB stained samples were homogenized in perchloric acid and after centrifugation the absorbance values for the supernatant were represented in the graph (**B**). Error bars represent the ± SD. Asterisks represent the significant difference in either fold level or percentage level in comparison with stress treated null segregants (NS) analyzed through ANOVA. * *P<* 0.05.

### OsANN5 induced Salinity stress tolerance is associated with altered SOD and catalase activity in recombinant *E.coli* and transgenic tobacco

Anti-oxidant enzymatic assay in *E. coli* revealed that the SOD activity was increased in the induced cultures of vector and *OsAnn5* transformed cells (Fig. 9A). However, heat and high salt treatment lead to decreased SOD activity in the same cultures. But, there was a further decrease in SOD activity in induced *OsAnn5* transformed cells as compared to induced vector transformed cells under stress treatments. In contrast to SOD activity, heat and high salt treatment lead to significantly increased CAT activity levels in the induced *OsAnn5* transformed cells in comparison to induced vector transformed cultures (Fig. 9B). Un-induced cultures were used as controls.

**Fig. 9.**
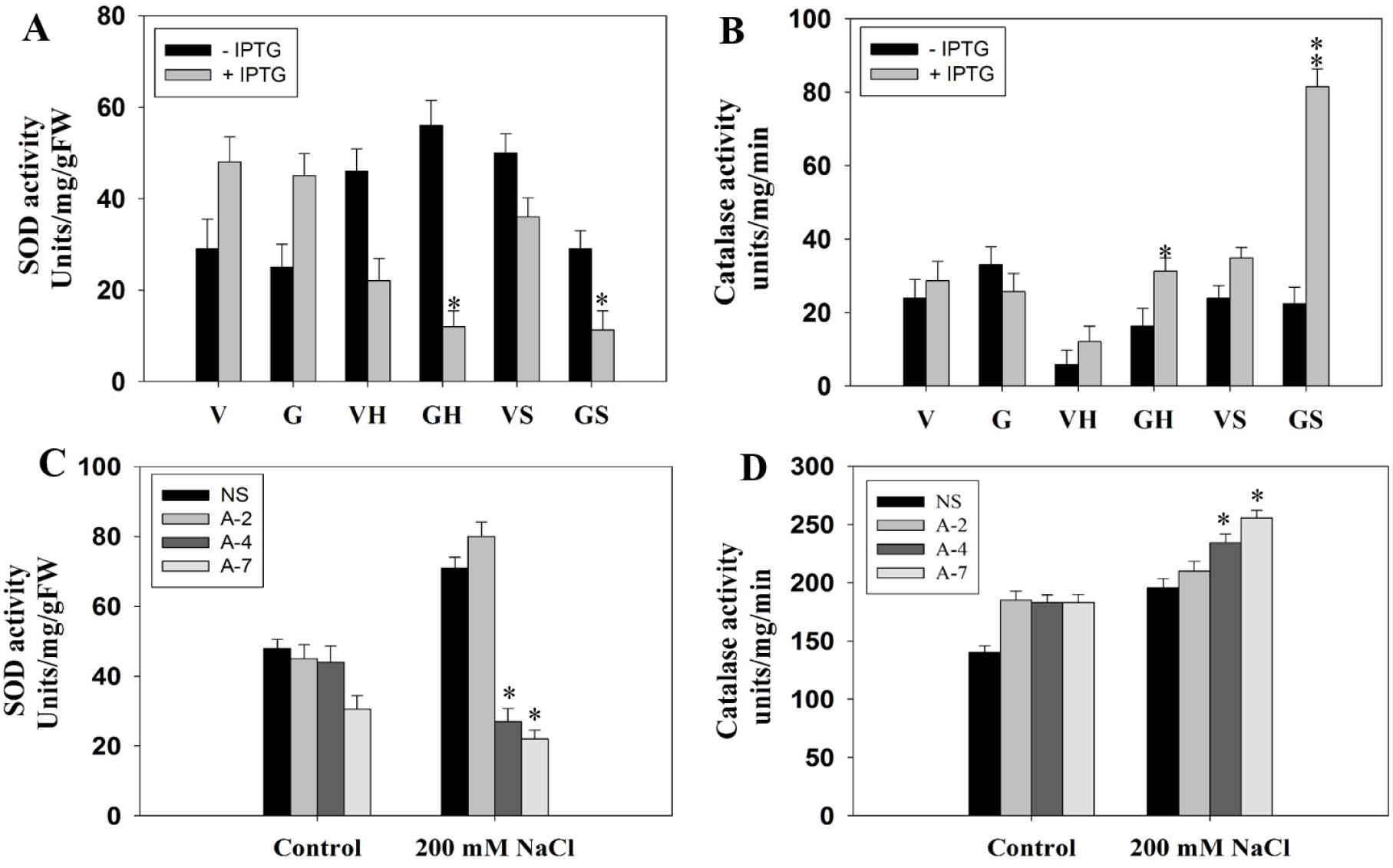
Quantification of anti-oxidant enzyme activity levels in the control and stress treated samples of *E.coli* and transgenics tobacco. In *E.coli* cells, SOD activity **(A)** and CAT activity **(B)** were determined for the pET32a vector (V) and pET32a:*OsAnn5* (G) transformed cells with IPTG (+ IPTG) or without IPTG (- IPTG) induction under heat stress (VH or GH= Heat stress given V or G samples) and high salinity stress (VS or GS = salt stress given V or G sample). In tobacco the SOD activity **(C)** and CAT activity **(D)** levels were measured for null segregant and 35S:*OsAnn5* transgenics samples under control and 200 mM NaCl treatments. Error bars represent the ± SD. Asterisks represent the significant difference in the activity levels of SOD/ CAT in comparison of VH with GH and VS with VH samples in the panel A and B, in comparison to the salt stress treated NS samples of tobacco in panel C & D when analyzed through ANOVA. * *P<* 0.05, * * *P<* 0.01.

Interestingly the 35S:*OsAnn5* tobacco transgenics also showed significant reduction in SOD activity (Fig. 9C) and increased CAT enzymatic activity (Fig. 9D) under the high salt stress compared to the control samples. The above results suggests that OsANN5 mediated oxidative stress tolerance is through the regulation of antioxidant enzymes.

### ABA-independent expression of *OsAnn5* under salt stress

To check whether the *OsAnn5* expression is independent of or dependent on ABA under salt stress, rice seedlings were treated with ABA, salt, and a combination of salt and fluridone and the transcript levels of *OsANN5* were quantified by qRT-PCR. The *OsAnn5* transcript levels were quantified along with the reference genes. It was observed that *OsAnn5* transcripts were doubled after 6 h of ABA treatment suggesting that *OsAnn5* responded to ABA. A significant eight fold increase was observed after 24 h of 200 mM NaCl salt stress treatment. But, the same salt treatment in combination with fluridone showed an approximately seven folds upregulation of *OsAnn5* transcript levels. This suggests that the fluridone treatment did not significantly affect the *osAnn5* expression under the salt stress. When we compared the basal expression of *OsAnn5* under normal and fluridone treatments, no significant change in the expression levels was observed. This suggests that treatment with fluridone did not affect the *OsAnn5* expression *in vivo* (Fig. 10A).

**Fig. 10.**
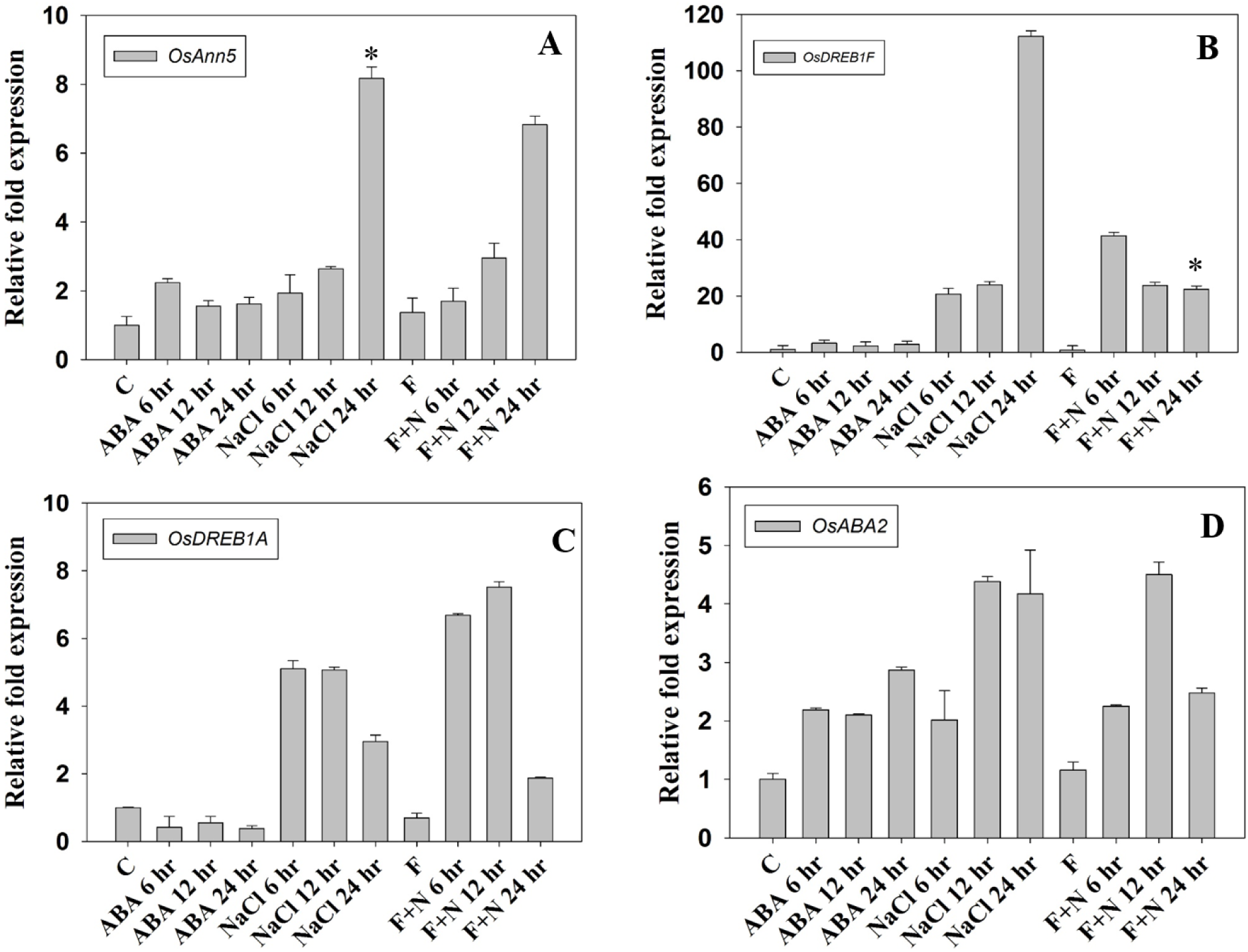
Real-time PCR quantification of *OsAnn5* **(A)***, OsDREB1F* **(B)***, OsDREB1A* **(C)***, OsABA2* (**D**) genes at different time points of the control, ABA (100 µM), NaCl (0.2 M) and fluridone (100 µM) (A) treatments. Error bars represent the ± SD. Asterisks represent the significant difference in either folds level or percentage level in comparison with control samples in panel A and with NaCl 24 hr treated samples in panel B when analyzed through ANOVA. * *P<* 0.05.

To check whether fluridone treatment inhibited the synthesis of ABA, we studied the expression levels of some genes whose expression under ABA has been determined earlier. Expression levels of an ABA-dependent gene *OsDREB1F* (Agarwal et al., 2010) was reduced from 112 folds to 22 folds at 24 h in the fluridone + salt stress treatment (Fig. 10B). There was no significant change in the expression levels of *OsDREB1A* gene, which is known to act through an ABA-independent pathway (Agarwal et al., 2010), (Fig. 10C). The transcript levels of *OsABA2* (ZEP) gene, which is known to be regulated by both ABA-dependent and independent pathway (Chen et al., 2014), did not change much during the 6 - 12 h of salt treatment. However, there was 1.8 folds decrease in the transcript levels under the combination treatment of fluridone with salt as compared with salt treatment at 24 h (Fig. 10D). These observations suggest that the expression of *OsAnn5* is independent of ABA during the initial stress treatment, with some dependence on ABA as the stress progressed.

In continuation with these observations, 10 d old *OsAnn5* transgenic and NS seedlings were grown on media supplemented with 4 µM and 8 µM ABA for 15 d (**Supplementary Fig. 5**). Their phenotype and their root length were observed after two weeks. Compare to NS, O*sAnn5* transgenic seedlings showed reduced sensitive to ABA (Fig. 11).

**Fig. 11.**
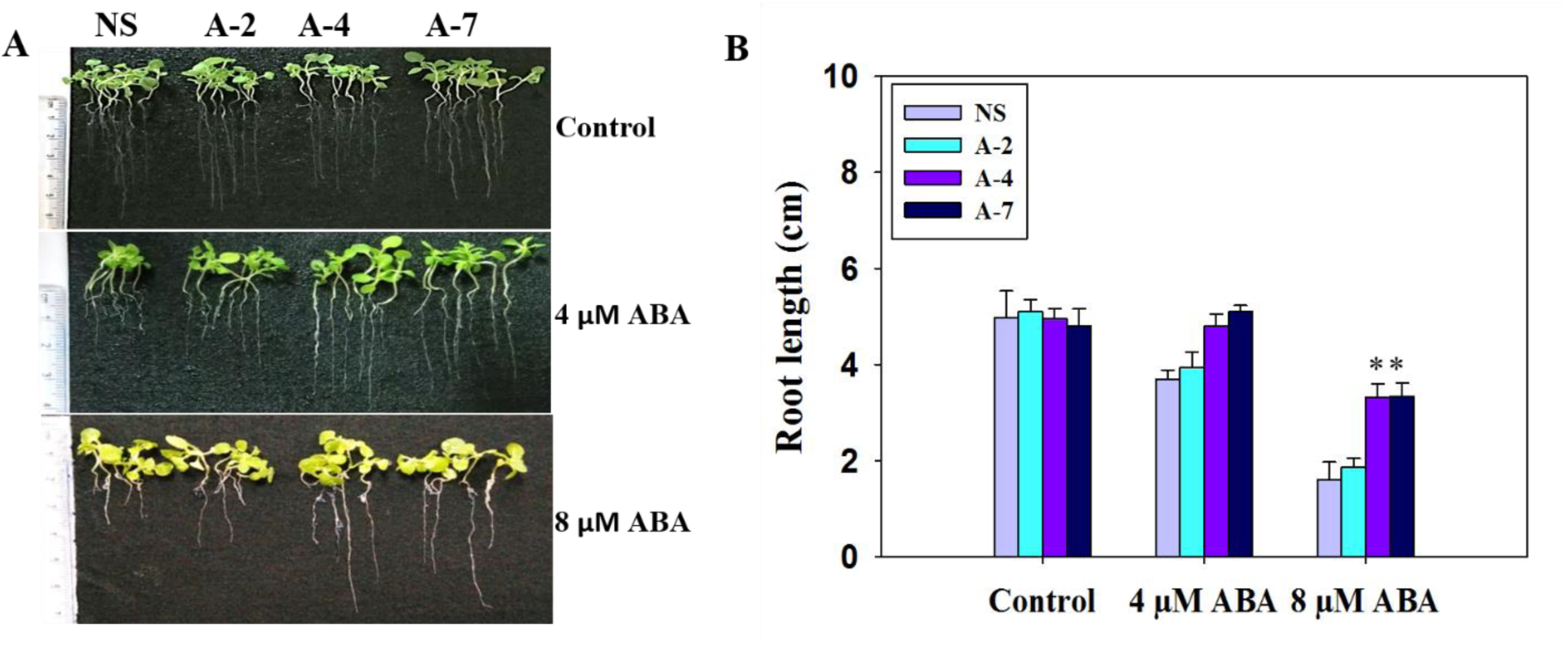
ABA sensitivity assay for control and 35S:*OsAnn5* transgenics at different concentrations of ABA **(A)**. After the treatment, differences in the root length was measured and represented in the graph. Error bars represent the ± SD. Asterisks represent the significant difference in either folds level or percentage level in comparison to null segregants under 8µM ABA treatment when analyzed through ANOVA. * *P<* 0.05

## Discussion

### *In silico* analysis for the upstream region of *OsAnn5* and mapping of important functional sites in OsANN5

A promoter defines when and where the transcription of a gene under its control should occur. Analysis of *cis*-elements in the promoter region generally gives an idea about its functionality that controls corresponding gene action. Various *cis*-elements related to abiotic stress conditions and hormones have been identified in the upstream region of *OsAnn5*. This suggests that these *cis*-elements are responsible for the involvement of OsANN5 in abiotic stresses and physiological responses.

ABRERATCAL is a Ca^+2^ responsive *cis*-element identified in the promoters of the 162 Ca^+2^ responsive genes (Kaplan et al., 2006). It was observed that one motif is enough to induce the overexpression of these genes. We found an ABRERATCAL *cis*-element in the *OsAnn5* putative promoter region. Apart from this a calcium binding motif was identified in the OsANN5 sequence. In support to this, in the calcium binding assays, OsANN5 showed calcium binding activity. Presence of calcium responsive *cis*-elements in the upstream region of *OsAnn5* suggests, Ca^+2^ ions not only regulates the *OsANN5* function but, also its expression. We also observed the presence of potential N-glycosylation and ubiquitination sites near to the calcium binding motif. This leads to a speculation that the calcium binding activity may be regulated by the PTMs.

The *OsANN5* protein was identified as a probable interacting partner of MYB through the STRING software. In earlier study, Punwani *et al*., 2007 observed that *Arabidopsis thaliana Annexin8* (*AtANN8*) expression by MYB98 transcription factor in synergid cells. This indicates that member of the MYB transcription factor family could be a possible upstream regulator of *OsAnn5*, which needs further experimental verification.

The possible occurrence of post-translational modifications in *OsAnn5* can be another level of regulatory mechanism, which can alter protein function by altering cellular and sub-cellular location, protein interactions and the biochemical reaction chains. If a gene is translated into a protein, its functional activity will be majorly directed by the posttranslational modifications. There are few reports on the PTMs of plant annexins (Lindermayr et al., 2005; Gorecka et al., 2005; Konopka-Postupolska et al., 2009). It was observed that posttranslational modification can alter the functional activity of the annexins (Konopka-Postupolska et al., 2009; Gorecka et al., 2005). Phosphorylation, glycosylation, acetylation, ubiquitination and SUMOylation are identified as potential posttranslational modifications that can modify the OsANN5 protein.

We found some highly potential phosphorylation and N-Glycosylation sites in OsANN5, which strongly suggest its functional importance in perceiving or transducing the signals. In our study, we identified the overlapping of SUMO interaction site and salt bridge motifs. This overlapping might have a function related to salt bridge formation in OsANN5. *In silico* prediction of potential PTMs can be explored for further experimental verification. This suggests a need for accelerated research towards post-translational modification of plant annexins for defining and clearly understanding the molecular mechanism of annexins.

Disordered regions in the protein are dynamic in nature and have the tendency to interact with other proteins or are prone to PTMs. The existence of disordered sequence with potential phosphorylation and O-glycosylation sites in the N-terminal region of OsANN5 suggests that it might play a major role in the signaling process in various environmental conditions. Crystal structure analysis of bell pepper annexin (AnxCa32) revealed that its core region interacts with the short N-terminal region (Hofmann et al., 2000a). Similarly, N-terminal and core disorder regions of OsANN5 possibly interact with each other under stress conditions through their disordered regions.

The alignment of OsANN5 with other plant annexins revealed an unconserved stretch of seven Glycine (Gly) residues. A stretch of Gly residues may decide the localization of the corresponding protein. Poly-glycine stretch in toc75 was necessary for targeting it to chloroplast envelope. Replacement of this glycine stretch with alanine made it miss-targeted to stroma (Inoue et al., 2003). Hence, it can be predicted that Gly stretch of OsANN5 may have a functional role in its localization.

### OsANN5 is localized in the peripheral regions of the cell after salt treatment

A given annexin can occupy various positions in cells, which include apoplast, organelles, and in association with membrane (Clark et al., 2012). Tissue-specific and sub-cellular localization of a specific protein may help in the identification of its function. As Ca^2+^ dependent membrane-binding proteins, many annexins possess the ability to dynamically change their cellular localization during certain physiological responses. For example, AtANN1 can be cytosolic, extracellular or can be associated with the plasma membrane, tonoplast and organelle membranes indicating that it can act as a multi-functional protein (Clark et al., 2010). The association of VCaB42 with vacuoles indicates its possible role in early vacuolar biogenesis (Seals et al., 1997). ANNSp2 was localized in the nucleus (Ijaz et al., 2017) predicting that it may have a transcriptional regulatory role. The fluorescence imaging studies reveals that OsANN5 is localized in the corners of the cells under normal condition. The plasmolysis treatment suggests that OsANN5 is not localizing to the cell wall. The salinity stress leads to OsAnn5 accumulation either in cytosol or plasma membrane which need further investigation.

### *OsAnn5* ameliorate stress tolerance through modulation of the anti-oxidant enzymes

*E. coli,* being a simple prokaryotic system, is the first choice for the heterologous expression of genes. There are many studies, which tried to analyse the functional role of the eukaryotic proteins in the *E*. *coli* system (Zhou et al., 2014; Kumari et al., 2009; Hu et al., 2014). Gidrol et al. (1996) observed that an annexin like protein rescued the ΔoxyR mutant of *E. coli* from the H_2_O_2_ stress. Similarly, the purified AtANN1 from *E. coli* cells exhibited the peroxidase activity, despite the possibility of the lack of posttranslational modification in the prokaryotic system. But, the same *ANNAt1,* purified from the *N. benthamiana* exhibited three times higher peroxidase activity (Gorecka et al., 2005) indicates that PTMs might have enhanced its peroxidase activity.

In the present study we also tried to assess the functional activity of the OsANN5 under different abiotic stresses by overexpressing it in *E. coli* cells. OsANN5 overexpressing *E. coli* cells were able to resist the given high salt, heat and drought stress conditions suggesting, that OsANN5 functions through a signaling pathway, which is common to both prokaryotes and eukaryotes in terms of the anti-oxidant systems.

From previous study It is already known that *OsAnn5* shown very quick response to heat stress (Jami et al., 2012). Spot assay also revealed that OsANN5 over expressing *E*.*coli* cells were also able to recovered even after 1 h of heat treatment. This results indicates that *OsAnn5* regulates the thermal stability of cells under stress condition.

To understand the functional role of *OsAnn5* in the abiotic stress, *OsAnn5* tobacco transgenics were developed and various stress assays were conducted. The ectopic expression of *OsAnn5* increased the stress tolerance of tobacco at seed germination stage, seedling stage and at mature plants level to the abiotic stresses in comparison to NS control. Assays with the seedling and leaf discs showed similar results in different abiotic stress treatments. This suggests that *OsAnn5* overexpression provides stress tolerance at different stages during tobacco plant development. At the seedling stage, NS control and low expression lines exhibited significantly reduced root length and stunted growth under stress conditions compared to the high expression lines. Biophysical analysis revealed that the stress tolerance of 35S:*OsAnn5* transgenics are correlated with enhanced osmolyte (proline) accumulation, reduced peroxidation and chlorophyll content in the cells. This kind of correlation was also observed with other annexin transgenics (Ahmed et al., 2017; Huh et al., 2010). DAB staining assay results supported the MDA quantification results for 35S:*OsAnn5* transgenics under salt stress.

The stress tolerance is generally associated with increased antioxidant defense system of the plant (Wang et al., 2003). It is observed that the anti-oxidant enzyme activity levels vary from plant to plant, kind and intensity of the stress and age of the plants (Mýtinová et al., 2010). In our study, we observed increased CAT activity and decreased SOD activity under the salt or heat stress treatments in both *OsAnn5* transformed *E.coli* and *OsAnn5* tobacco transgenics. These results strongly suggest that *OsAnn5* modulates CAT and SOD levels during the stress conditions and attributes to the stress tolerance of OsANN5. This also suggests that *OsAnn5* could act as a balancing protein by inducing the ROS mediated signaling and scavenging the ROS.

### *OsAnn5* expression is possibly independent of ABA under salt stress *in vivo*

Many of the plant annexins are well known for their crucial role in biotic and abiotic stress tolerance and developmental processes (Yadav et al., 2018). But, the signaling pathways in which they are involved are yet to be clearly identified. Abscisic acid is a key signaling intermediate during abiotic stress, which plays a crucial role in developmental and drought stress tolerance.

Many genes would be upregulated or downregulated by this hormone. Some recent reports tried to elucidate the role of annexins in correlation with ABA in stress signaling pathway. For example, Bianchi et al. (2002) suggested that AnnAt1 could be acting downstream of a cross talk between ABA-Auxin. A *Solanum pennellii* annexin, Annsp2 is associated with ABA accumulation during drought stress when over expressed in tomato and showed insensitivity to external ABA application during the seed germination stage (Ijaz et al., 2017). The annexin, *AnnBj2* overexpressing transgenic plants of mustard were also insensitive to ABA during the seed germination stage. Interestingly, gene expression studies in the *AnnBj2* transgenic mustard plants revealed increased transcript levels of ABA catabolic gene *CYP707A2* (Ahmed et al., 2017). In line with these results, *OsAnn3* knock down transgenics showed ABA sensitivity at seed germination stage (Shen et al., 2017) suggesting that annexins may enhance seed germination by promoting the degradation of ABA. The above studies strongly suggest the functional role of annexins in abiotic stress tolerance that is associated with ABA. Hence, as a first step towards understanding OsANN5 involvement in ABA signalling pathway, we tried to assess whether any alteration in ABA levels would affect *OsAnn5* expression in stress condition, which eventually gives a clue asto whether it is ABA-dependent or independent or both.

Earlier jami et al., reported that no rice annexin got responded to the ABA treatment. In our case *OsAnn5* shown upregulation at 6 h of ABA treatment (**Supplementary Fig. 6**). So we tried to know whether *OsAnn5* expression dependent of ABA under salt stress. We observed not much significant difference in the expression levels of the *OsAnn5* when compare with salt stress and salt stress in combination with fluridone. In line with this, 35S:*OsAnn5* transgenic seedlings showed reduced sensitivity to external ABA. This clearly suggests that OsAnn5 may acts majorly through an ABA independent pathway under salt stress. The presence of ABRE along with DRE in the *OsAnn5* upstream region suggesting *OsAnn5* possibly act through ABA-dependent and independent pathway under stress conditions.

## Conclusion

Presence of several probable post translational modifications sites in annexins suggesting that it can play a major role in stress tolerance. This was evinced through heterologous overexpression of *OsAnn5* in *E.coli* and tobacco, which resulted in their enhanced tolerance to abiotic stresses. Under salt stress, *OsAnn5* works through an ABA-independent pathway. ROS scavenging activity exhibited by 35S:OsAnn5 transgenics was correlated with the modulation of their antioxidant enzyme levels elucidating the possible involvement of OsAnn5 in stress tolerance. Hence, a new dimension of research related to the identification of the binding partners and post-translational modifications of OsAnn5 would throw more light on the detailed mechanism of its action.

## Author contribution

PB and PBK planned and executed the work and TS, DY and HK assisted in analyzing the data and writing the manuscript.

## Acknowledgements

The authors are grateful to the Head, Department of Plant Sciences for facilities under DST-FIST and UGC-SAP. PB and TS acknowledge Ms. V. Aswani for her timely help during the experiments.

