## supplemental information for "Ectopic overexpression of abiotic stress-induced rice annexin, *OsAnn5* potentiates tolerance to abiotic stresses"

**Supplementary Material**

**Ectopic overexpression of abiotic stress-induced rice annexin, *OsAnn5* potentiates tolerance to salt, drought and osmotic stresses**

Prasanna Boyidi*, Trishla Vikas Shalibhadra, Halidev Krishna Botta, Deepanker Yadav, Pulugurtha Bharadwaja Kirti ^⃰^

Department of Plant Sciences, School of Life Sciences, University of Hyderabad, Prof. C. R. Rao Road, Gachibowli, Hyderabad, 500046

*Corresponding authors at Department of Plant Sciences, School of Life Sciences, University of Hyderabad, Prof. C. R. Rao Road, Gachibowli, Hyderabad, 500046, India.

**Supplementary Table 1:** List of primers used in the study

**Primers for *OsAnn5* amplification**

OsAnn5 FP: 5’TAT GGT ACCATG GCG AGC CTG AGC GT’3 (KpnI)

OsAnn5 RP: 5’ATT GGA TCCTTA GCG GTC GCG GCC’3 (BamHI)

**Primers used to clone *OsAnn5* into pEGAD vector**

OsAnn5 FP: 5’TAT GAA TTC ATG GCG AGC CTG AGC GT’3 (EcoRI)

OsAnn5 RP: 5’ GTA AGC TTT TAG CGG TCG CGG C ’3 (HindIII

**Supplementary Table 2:** Various bio-informatics related web tools used in the analysis of upstream promoter region of *OsAnn5* and its deduced amino acid sequence

| **1** | **http://www.dna.affrc.go.jp/PLACE/** | ***OsAnn5* upstream region analysis tool** |
| --- | --- | --- |
| **2** | **http://www.cbs.dtu.dk/services/NetPhos/** | **Phosphorylation prediction tool** |
| **3** | [**http://bdmpail.biocuckoo.org**](http://bdmpail.biocuckoo.org) | **Acetylation site prediction tool** |
| **4** | [**http://sumosp.biocuckoo.org/online.php**](http://sumosp.biocuckoo.org/online.php) | **Sumoylation prediction tool** |
| **5** | [**http://bdmpub.biocuckoo.org/**](http://bdmpub.biocuckoo.org/) | **Ubiquitination prediction tool** |
| **6** | [**http://crdd.osdd.net/raghava/glycoep/index.html**](http://crdd.osdd.net/raghava/glycoep/index.html) | **Glycosylation sites prediction tool** |
| **7** | [**https://string-db.org/**](https://string-db.org/)**version 10.5** | **Interacting partners prediction tool** |

**Supplementary Table 3**: List of putative *cis*-elements and transcription factor binding sites identified in the 1.5 kb upstream region of the *OsAnn5.*

| **S.No** | **Cis-elements Name** | **Sequence** | | **No** | **Function** |
| --- | --- | --- | --- | --- | --- |
| **Cis-elements involves in Abiotic stress** | | | | | |
| 1 | DRECRTCOREAT | | RCCGAC | 2 | Dehydration responsive elements |
| 2 | ACGTATERD1 | | ACGT | 18 |  |
| 3 | CBFHV | | RYCGAC | 2 |  |
| 4 | LTRECOREATCOR15 | | CCGAC | 3 | Low temperature, Drought response elements |
| 5 | CCAATBOX | | CCAAT | 1 | Responds to Heat |
| 6 | GT1GMSCAM4 | | GAAAAA | 2 | Salt inducible |
| 7 | SURECOREATSULTR11 | | GAGAC | 2 | Sulfur responsive elements |
| 8 | CURECORECR | | GTAC | 4 | Copper response elements |
| 9 | ARE1 | | RGTGACNNNGC | 1 | Antioxidant response element |
| 10 | CGCGBOXAT | | VCGCGB | 20 | Ca^2+^/Calmodulin binding |
| 11 | WBOXATNPR1 | | TTGAC | 1 | WRKY transcription factor binding site |
| 12 | WBOXHVISO1 | | TGACT | 3 |  |
| 13 | WBOXNTERF3 | | TGACY | 4 |  |
| 14 | WRKY71OS | | TGAC | 15 |  |
|  | **Hormone response** | | | | |
| 16 | ASF1MOTIFCAMV | TGACG | | 7 | Auxin response elements |
| 17 | AUXRETGA2GMGH3 | TGACGTGGC | | 2 |  |
| 18 | SEBFCONSSTPR10A | YTGTCWC | | 1 | Similar to auxin response elements |
| 19 | ABREA2HVA1 | CCTACGTGGC | | 1 | ABA response elements |
| 20 | ABREATCONSENSUS | YACGTGGC | | 1 |  |
| 21 | ACGTABREMOTIFA2OSEM | ACGTGKC | | 3 |  |
| 22 | ABRELATERD1 | ACGTG | | 7 |  |
| 23 | ABRERATCAL | MACGYGB | | 1 |  |
| 24 | GARE1OSREP1 | TAACAGA | | 1 | Gibberellin response elements |
| 25 | TATCCACHVAL21 | TATCCAC | | 1 |  |
| 26 | ARR1AT | NGATT | | 10 | Induced by cytokine response genes |
| 27 | GCCCORE | GCCGCC | | 6 | Ethylene responsive elements |
|  | **Transcription factor binding sites** | | | | |
| 28 | MYBST1 | GGATA | | 4 | MYB binding elements |
| 29 | MYBCOREATCYCB1 | AACGG | | 1 |  |
| 30 | MYBCORE | CNGTTR | | 6 |  |
| 31 | TATCCAOSAMY | TATCCA | | 1 |  |
| 32 | MYB1AT | WAACCA | | 2 |  |
| 33 | MYB2CONSENSUSAT | YAACKG | | 1 |  |
| 34 | MYCATERD1 | CATGTG | | 1 | MYC binding elements |
| 35 | MYCCONSENSUSAT | CANNTG | | 12 |  |
| 36 | MYCATRD22 | CACATG | | 1 |  |
| 37 | DOFCOREZM | AAAG | | 7 | Dof transcription factor binding sites |
|  | **Elements for tissue specific expression** | | | | |
| 39 | CACTFTPPCA1 | YACT | | 12 | Mesophyll cells specific expression |
| 40 | POLLEN1LELAT52 | AGAAA | | 3 | Pollen specific expression |
| 41 | DPBCOREDCDC3 | ACACNNG | | 2 | Embryo specific and induced by ABA |
| 42 | OSE1ROOTNODULE | AAAGAT | | 2 | Organ specific elements usually present in the promoters of infected root nodules |
| 43 | OSE2ROOTNODULE | CTCTT | | 5 |  |
| 44 | RHERPATEXPA7 | KCACGW | | 4 | Root hair specific elements |
| 45 | ROOTMOTIFTAPOX1 | ATATT | | 1 | Root specific elements |
| 46 | RAV1AAT | CAACA | | 3 |  |
| 47 | IBOXCORE | GATAA | | 11 | Light responsive elements |
| 48 | SORLIP2AT | GGGCC | | 7 |  |
| 49 | GT1CONSENSUS | GRWAAW | | 5 |  |
| 50 | ASF1MOTIFCAMV | TGACG | | 7 |  |
| 51 | SORLIP1AT | GCCAC | | 13 |  |

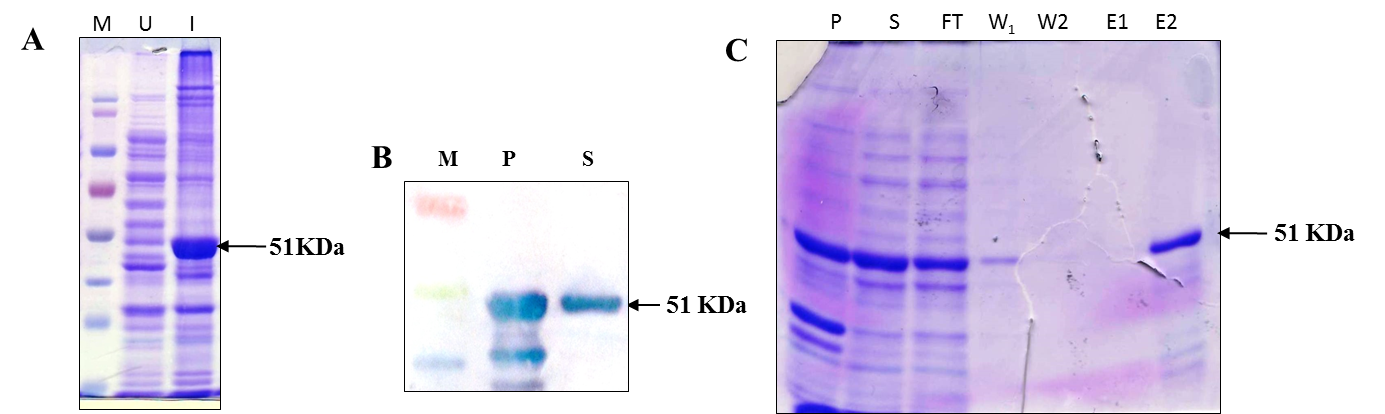

**Supplementary Fig.1** Confirmation of the overexpressed OsAnn5 protein using SDS-PAGE A. M= marker, U = Un induced cells, I = Induced cells. B. Western blot confirmation of OsANN5 from the transformed cells, P= Pellet, S= Supernatant. C. OsAnn5 protein purification through Ni-NTA column using His tag (C), P= Pellet, S= Supernatant, FT= Flow through, W1= Wash1 (20mM Imidazole, W2= wash2 (30mM Imidazole), E1= Elution1(100mM Imidazole), E2= Elution2 (200mM Imidazole)

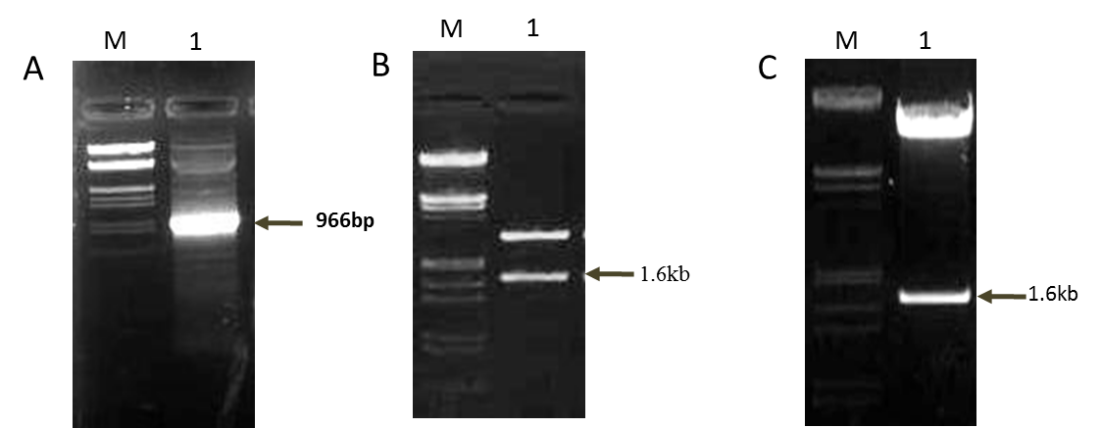

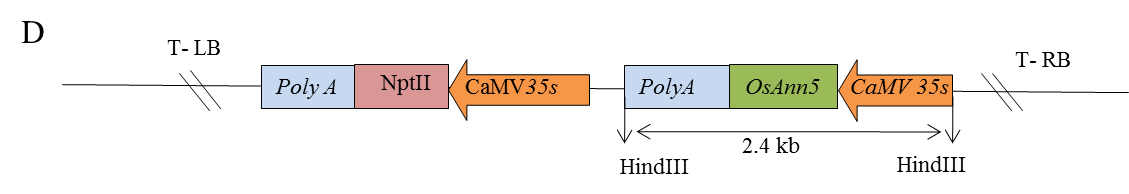

**Supplementary Fig. 2** Development of the recombinant binary vector, pCAMBIA2300:CaMV35S:*OsAnn5* (**A**). Confirmation 966 bp *OsAnn5* CDS was PCR amplified from the rice total cDNA, B. Confirmation of OsAnn5 expression cassette from pRT100, C. Confirmation of the final 35S:*OsAnn5:poly*A that was cloned into pCAMBIA2300 vector. Pictorial representation of T-DNA region of pCAMBIA2300 harbouring OsAn5 cassette along with *nptII* as selection marker (**D**).

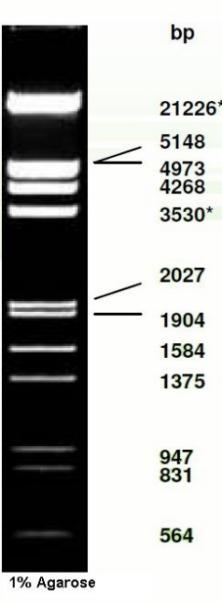

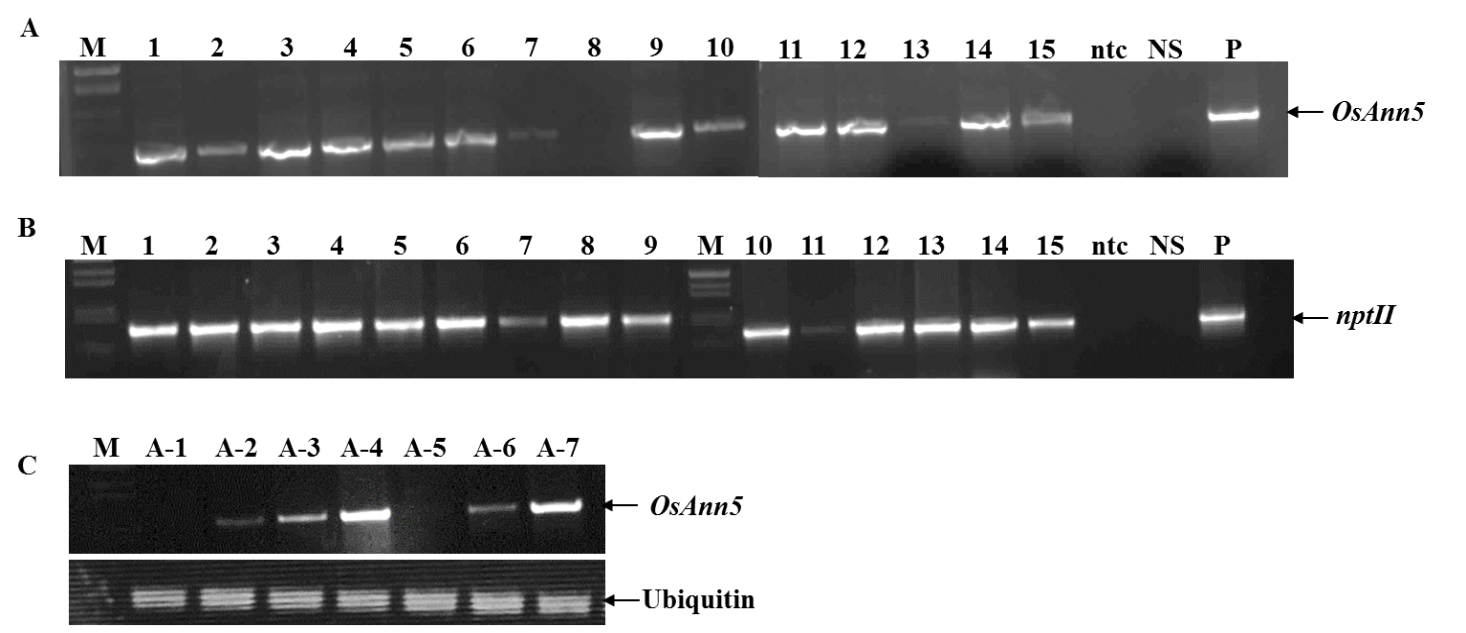

**Supplementary Fig. 3** PCR confirmation for the 35S:*OsAnn5* transgenics using their genomic DNA for the presence of 966 bp *OsAnn5* (**A**) and 720 bp *nptII* (**B**). Semi-quantitative PCR for the identification of low and high expression lines (**C**) along with the ubiquitin as a reference gene. M= Lambda *Eco*RI and *Hin*dIII Marker, 1-15 independent primary transgenic plants, NS= Null segregant, ntc = non templet control, P= Positive control

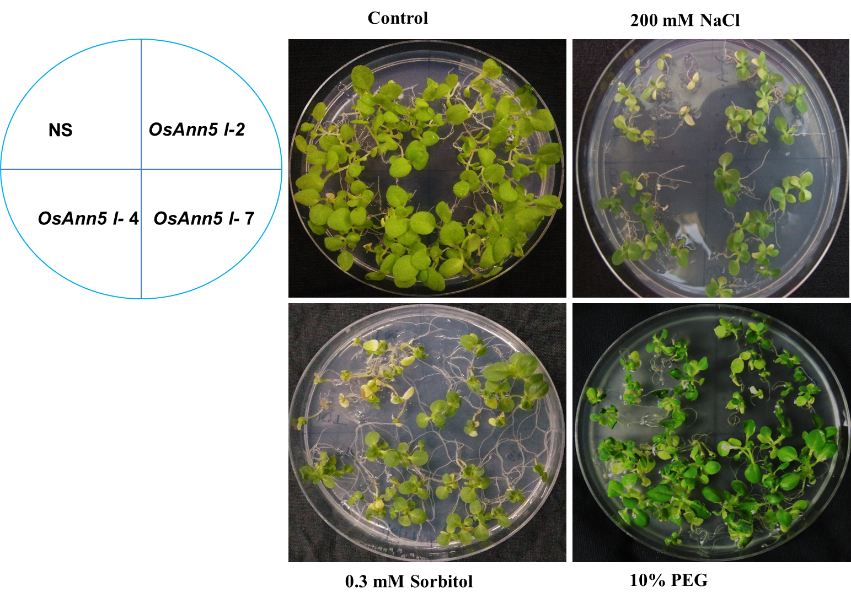

**Supplementary Fig. 4** Compared to the NS, 35S:*OsAnn5* transgenics were able to tolerate different abiotic stresses like 200 mM NaCl, 0.3 M Sorbitol, 10% PEG.

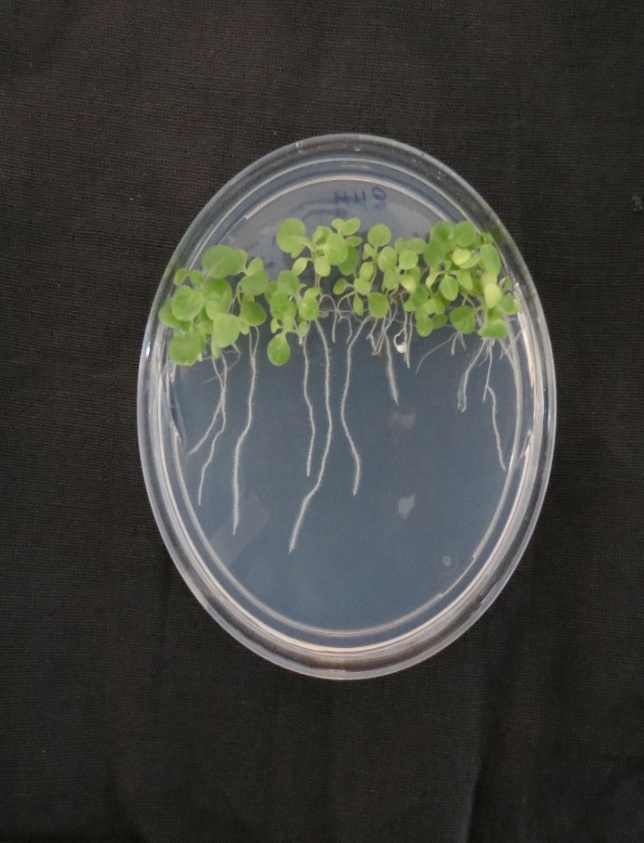

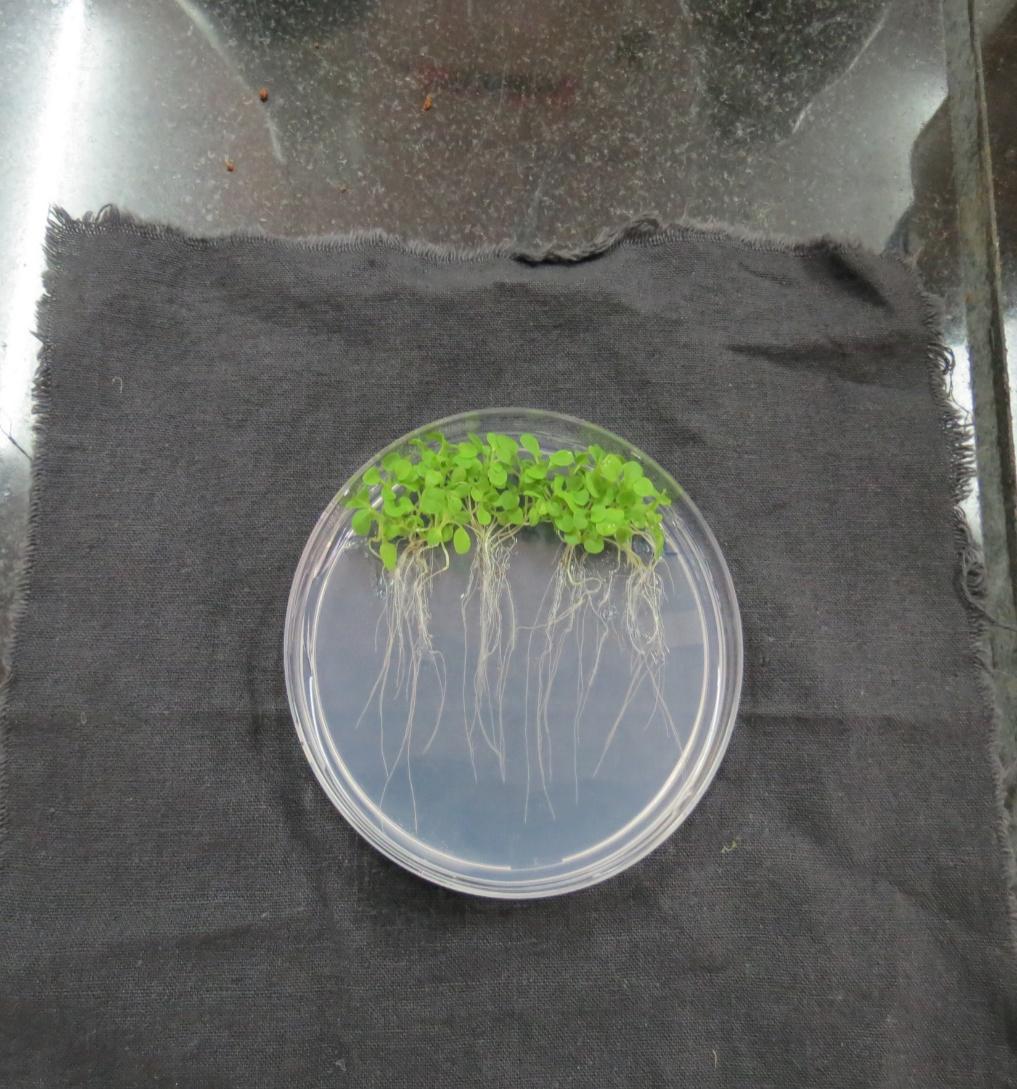

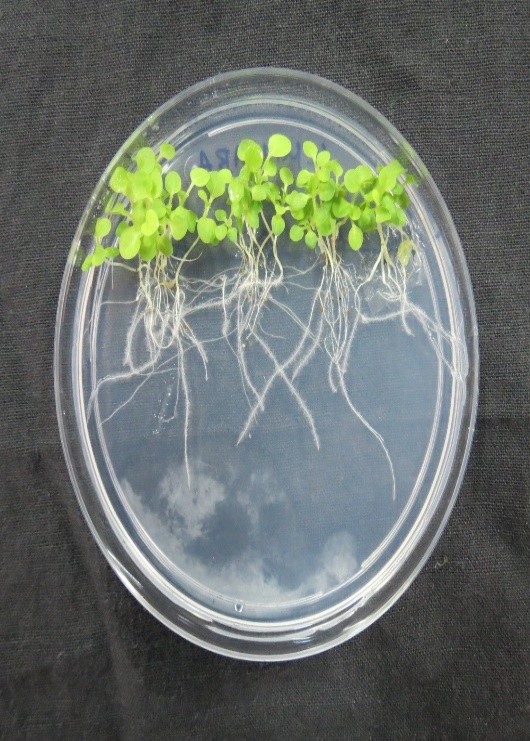

**Control	4 µM ABA	8 µM ABA**

NS	A-2	A-4	A-7	NS	A-2	A-4	A-7	NS	A-2	A-4	A-7

**Supplementary Fig.5** 35S:*OsAnn5* transgenics showing insensitivity to external application of ABA compared to the null segregant (NS)

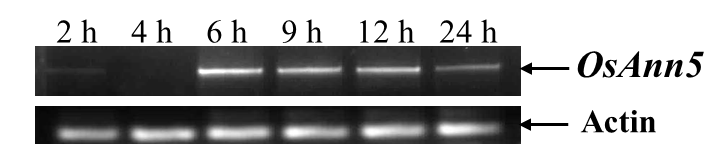

**Supplementary Fig. 6** Semi-quantitative PCR for the ABA (100 µM) treated 10 d rice seed lings with reference to the actin gene expression. The samples were collected at different time points during the treatment (2 h, 4 h, 6 h, 9 h, 12 h, 24 h).
